# Climate Shaped the Global Population Structure of Leopards and their Extinction in Europe

**DOI:** 10.1101/2025.10.03.680393

**Authors:** Zhe Xue, Johanna L.A. Paijmans, Andrea Vittorio Pozzi, Sidney Leedham, Michela Leonardi, Cecilia Padilla-Iglesias, Margherita Colucci, Anahit Hovhannisyan, Andrea Manica

## Abstract

Environmental change is often invoked as a key force shaping species evolution and demography, but quantifying its role is challenging. Leopards (*Panthera pardus*), a widely distributed generalist species, provide an ideal case for studying the role of the environment. The population dynamics of African and Asian leopards differ dramatically, with near panmixia in Africa versus a strong structure and eight subspecies in Asia. Fossil records show that a population in Europe disappeared after the Last Glacial Maximum (LGM), further pointing to complex range dynamics in Eurasia. In this study, we explicitly test the role of climate in shaping leopards population dynamics across the continents, by quantitatively combining paleoclimatic, demographic, and genetic simulations over the last 450 thousand years. Using an Approximate Bayesian Computational framework, we show that the genetic structure differences between Africa and Asia can be explained by distinct historical climatic conditions in these two continents: most of sub-Saharan Africa and parts of Southeast Asia exhibited a stable range without geographical barriers, while other areas such as Morocco, Afghanistan and Northeast Asia showed expansion and extinction cycles during glacial and interglacials. We further model population dynamics in Europe and validate it against fossil and radiocarbon dates records. We find that European populations were likely fragile, with extinctions repeatedly predicted over multiple glacial cycles, indicating that climate change possibly led to the extinction of the European subspecies following the LGM. Notably, we found that climate during the Holocene should have allowed a more recent recolonisation that did not happen, indicating other factors such as human presence might have blocked it.

## Introduction

Environmental change is often invoked as a major driver of population differentiation, shaping the structure of populations and species worldwide. However, quantitatively testing the role of such change in shaping differentiation is challenging. This lack of quantification is not only a conceptual issue, but it also limits our ability to forecast the role of contemporary human-induced global changes on biodiversity. Thus, reconstructing how past climate has shaped species’ evolutionary and demographic dynamics has both scientific as well as practical relevance.

Leopards (*Panthera pardus*) provide an ideal system to study the role of the environment in shaping population demographics and the initial process of differentiation. Population genetics studies have shown dramatic differences in population dynamics across the species’ broad range, shaped after an out-of-Africa event approximately 300 - 600 thousand years ago estimated with genomic data (Paijmans et al., 2018, Paijmans et al., 2021; Pecnerova et al., 2021;). The single subspecies found in Africa has shown weak structure and strong mobility, typical for a far-ranging large predator adapted to different habitats, while its Asian range is highly structured, showing signs of strong isolation by distance (Paijmans et al., 2021; Pecnerova et al., 2021), and is currently split into eight subspecies by the IUCN (Stein et al. 2013), although an alternative taxonomy of seven Asian subspecies has been proposed (Kitchener et al. 2017). Despite these advances, it is still unclear what shaped such highly contrasting differentiation patterns in the two continents.

Climate is known to be a major driver for fauna evolutionary changes in many cases. For leopards, their environmental niche spans a wide range of habitats on both continents, with little sign of Asian leopards having expanded beyond the niche observed in Africa except Persian leopard (Leedham et al., 2023). Thus, the differentiation observed in Asia is possibly not the consequence of local adaptation to novel environments after their expansion from the ancestral African range. It is plausible that geographic barriers, coupled with greater climatic instability in the Asian continent compared to their ancestral African home, have driven complex dynamics of colonization and extinction.

The role of climate in determining the extinction of the European population at the end of the Last Glacial Maximum (LGM) is another highly debated question. The fossil record suggests that leopards once inhabited a wide area spanning from southern to central Europe, with modern-leopard-like fossils dating back to the middle Pleistocene and lasting until the LGM (Diedrich 2013), during which the occurrences in central Europe disappear. It is also unclear whether this disappearance, like many other megafauna extinctions observed in the same period, was primarily driven by climate, human activity, or a synergy between the two (Bergman et al., 2023; Prescott et al., 2012), as shown for the disappearance of European bison (Pilowsky et al., 2023).

To test the influence of climate on the population dynamics and genetic structure of leopards, multidisciplinary approaches that quantitatively integrate multiple lines of evidence are needed. Climate-Informed Spatial Genetic Models (CISGeM) are frameworks that aim to use paleoclimatic models to inform demographic and genetic simulations of a species based on the environmental change through time and space. The parameters that best describe the past demography of a species can be formally chosen by comparing the predicted patterns of differentiation among samples with the observed quantities from genetic data using an Approximate Bayesian Computation framework.

These models have been successfully used to reconstruct the range expansion of Anatomically Modern Humans out of Africa (Eriksson et al., 2012; Delser et al., 2021), the shape of Pan-African metapopulation structure (Padilla-Iglesias et al., 2025), and to model how the contraction and expansion in the last glacial cycle shaped yellow warblers’ population structure (Miller et al., 2022).

Here, we use CISGeM to reconstruct the population dynamics of global leopards over the last 450 thousand years. We first calculated the genetic structure of worldwide leopards using modern and historical genomes, then fitted a model that successfully recaptured the structure between African and Asian leopards. The reconstructed demography showed that Africa and Asia were consistently separated by strong geographical barriers on the west bank of the Red Sea. Stable climate conditions for leopards in Africa shaped a panmictic population, while in Asia recurrent range fragmentation during glacial cycles created several strong bottlenecks that resulted in the observed differentiation between subspecies. Lastly, we found that leopards have likely gone extinct in Europe due to the unsuitable climate during LGM.

## Results and Discussions

### Spatio-temporal simulations successfully recover the genetic structure of leopards

We assembled a whole-genome dataset containing 15 samples from previous publications (Paijmans et al., 2021, Pecnerova et al., 2021, Kim et al., 2016) that represents global subspecies genetic structure, following the taxonomy suggestions proposed by Kitchener et al., 2017 (Fig. 1a). After processing the raw sequence data, accounting for DNA damage and differences in coverage in the modern and historical genomes, we quantified population structure as a matrix of pairwise differentiation (Fig 1d), using the mean of pairwise π estimated on 16,901 windows of 5,000 bp length (see methods). This pairwise matrix successfully captures the high level of differentiation in the genetic composition of leopards between the African and Asian continents, and their distinct isolation-by-distance pattern (Fig 1b, p-value_afr = 0.001, p-value_asia = 0.001 under 1,000 permutations) as previously reported by Paijmans et al. (2021)

**Figure 1.**
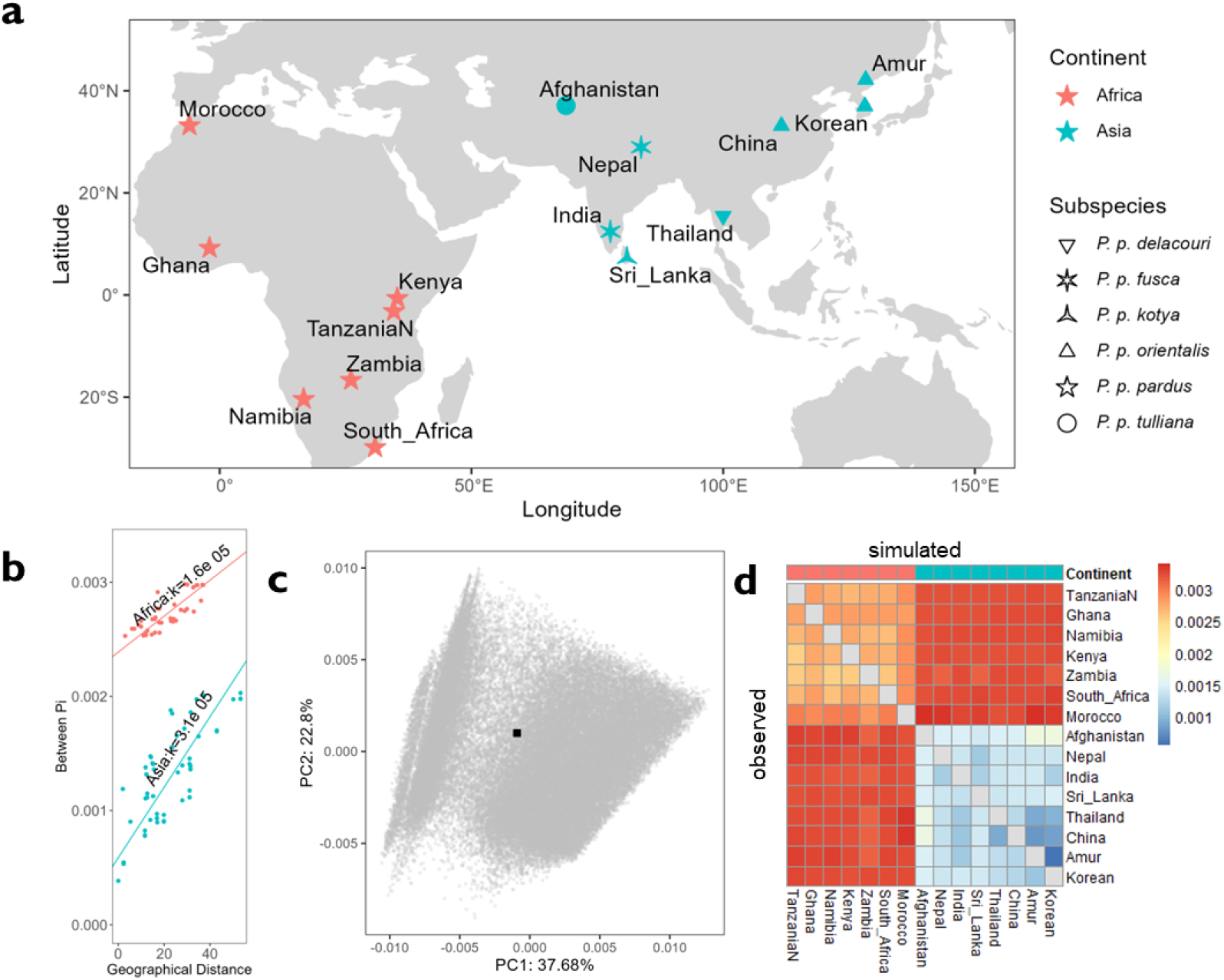
**a**, Geographic distribution and subspecies classification of the empirical sample genomes used in this study. **b**, The within continent isolation-by-distance, measured by between-group pairwise π vs geographical distance (Unit: 100 KM). The standard major axis (SMA) regression slopes for each linear regression is shown. **c**, The first 2 PCs of the summary statistics of simulations (grey points) and the empirical value (black point). **d**, The pairwise genetic distance matrix of the best simulations vs empirical. Lower Triangle: The empirical pairwise π between samples. Upper Triangle: Mean pairwise π of the top 1000 best simulations.

To investigate whether and how climate shaped the genetic structure of leopards, we used Species Distribution Modelling (SDM, Elith and Leathwick 2009) to reconstruct the suitability in Africa and Eurasia following Leedham et al., 2023. Briefly, based on 8,708 present and historical leopard occurrences data, we built a SDM using an ensemble approach (Araujo and New 2007), then used paleoclimate model outputs based on a statistical emulator of the HadCM3 global circulation model (Krapp et al., 2021) to project this ensemble into the last 450,000 years and create maps of the predicted past habitat ranges of leopards (Supplementary Figure 3-6). From these predictions, we then extracted the probability of occupation and projected them onto an equal-area hexagonal grid covering the whole of Africa and Eurasia as the background for CISGeM to define changes in suitability over time.

We then used CISGeM to perform spatial-temporal simulations of demography and genetic change based on these background suitability. In the model, for any given deme (i.e. a cell on the hexagonal grid), the carrying capacity was modelled as a function of the probability of occupation from the SDM. Individuals can freely migrate between adjacent cells, controlled by migration parameters. Thus, population sizes and migrations, and consequently the broader demography of leopards, were allowed to change over time, driven by climatic changes. During the spatial simulation the genealogies of loci carried by individuals are also tracked, allowing the measurement of various genetic summary statistics, e.g. pairwise π matrix between individuals, from simulated individuals in any demes. The posterior distribution of parameters that describe the link between probability of occupation, effective population size, migration and growth rate (See supplementary table for full parameters used in the simulations) were quantified using Approximate Bayesian Computation - Random Forests.(Raynal et al., 2019) on the matrix of pairwise π among all the genomes. (See methods section for detailed description of the method)

To explore the demographic history of leopards, we simulated 90,224 scenarios with random parameter combinations. We set the starting point in East Africa, as suggested by fossil records (Werdelin et al., 2010). For starting time, we selected 450,000 years BP as the starting point of simulation, an average time of the two previous whole-genome studies (500,000-600,000 BP from Paijmans et al., 2021; 300,000-400,000 BP from Pecnerova et al., 2021) that also approximately falls in previous mtDNA range (95% CI: 457,000 - 956,000, Paijmans et al., 2018). Among these simulations, 51,539 combinations of parameter values produced a demography that led to the expansion of all leopards samples into their present range, and genetics simulations among these samples were performed.

To validate the model, we then compared the simulated genetic differentiation pattern - pairwise π – against the observed ones. Each observed value of pairwise π fell within the range of the values produced by simulations (Supplementary Table 1.2). We further summarised the whole pairwise π matrix using Principal Component Analysis (PCA) to capture the relationships among pairwise π estimates and ensure that they were captured by the model, confirmed by that the observed PC scores for the empirical pairwise matrix were found within the cloud of points representing the simulations. (Fig. 1c)

To understand drivers and demographic process shaping the leopards genetics structure, we took the best 1,000 parameter combinations (defined as having the smallest Euclidean distances between observed and predicted pairwise π values), and used the mean of their resulting demographics as a representative aggregated demography that match the observed genetic pattern in modern leopards (Fig. 2 & 3; Supplementary Media 1). Overall, these simulated demography recreated the majority of distinct diversity patterns observed in Africa and Asia (Figure 1d), indicating that climate was a major driver of the global genetic structure found in leopards. African leopards kept a relatively stable range over several glacial cycles, leading to low levels of isolation by distance, whereas the demographic history in Asia is characterised by dynamic climate-driven expansions and contractions.

**Figure 2.**
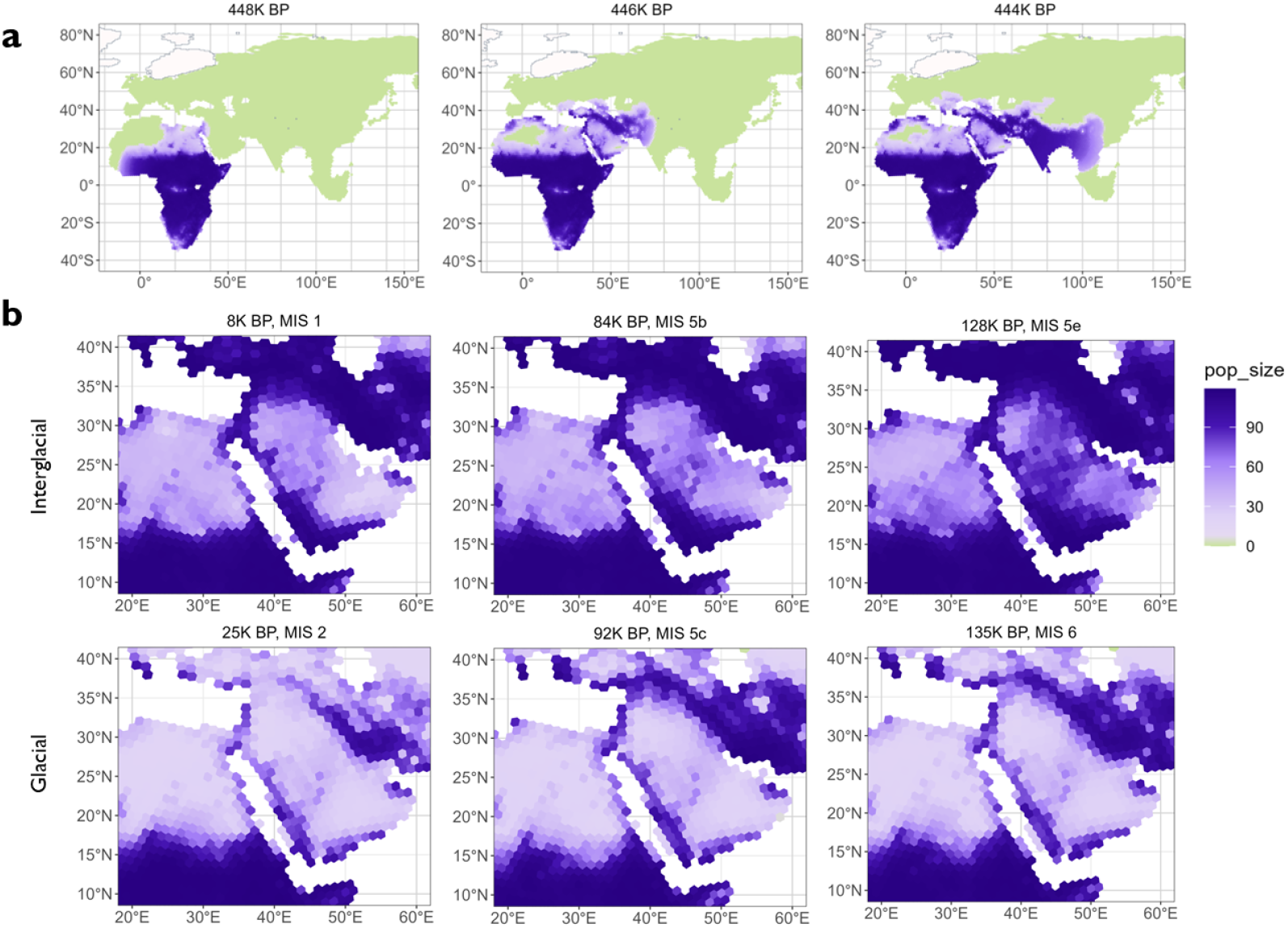
Fast Initial Expansion and Stable Inter-continental separation of leopards. A. Initial expansion as inferred by our model. B. The corridor of the west bank of the Red Sea during recent glacial cycles.

**Figure 3.**
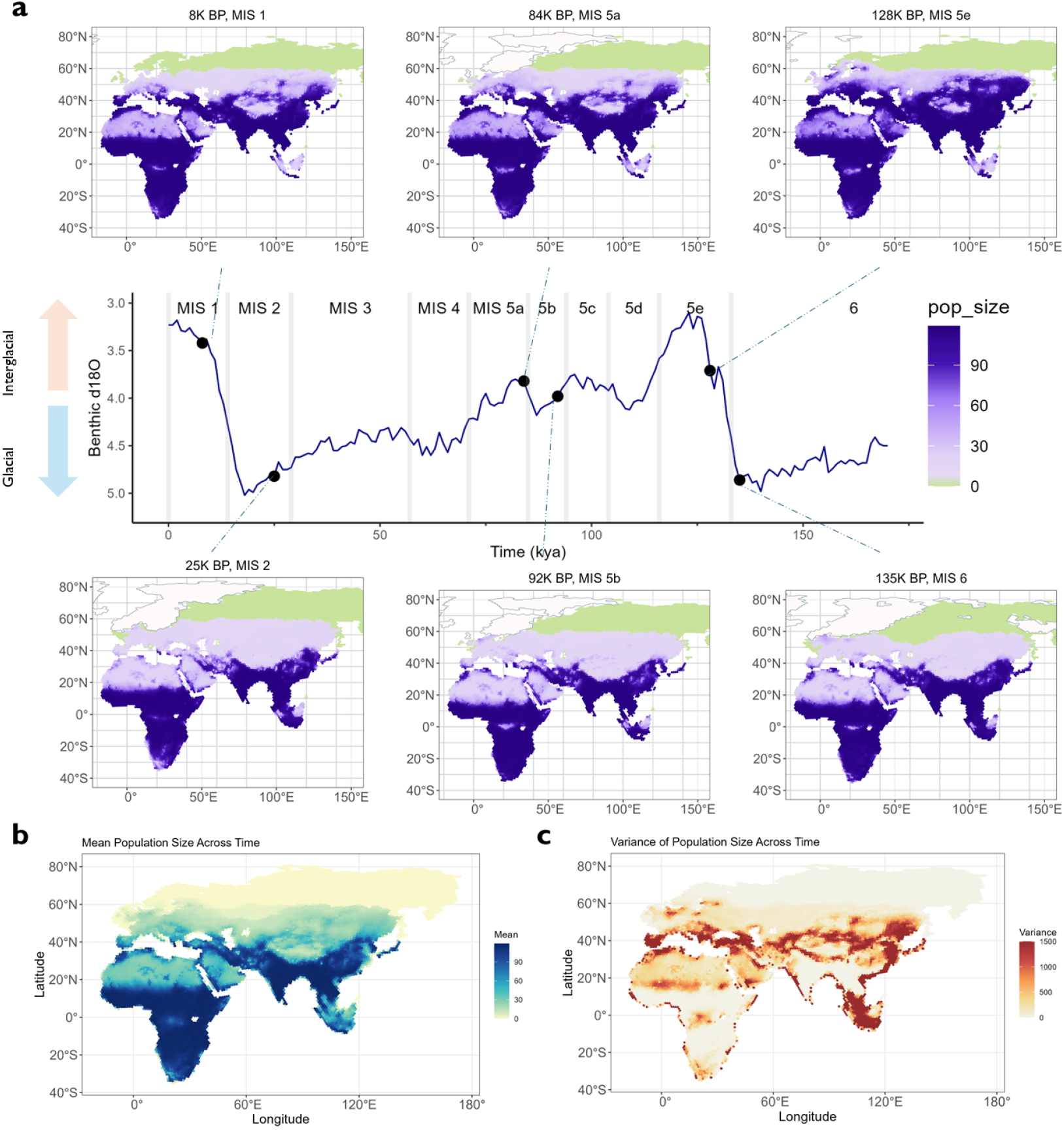
Worldwide population size fluctuation during recent glacial cycles. a, Mean population size inferred by the model at chosen timepoints during interglacial and glacial phases. White areas indicate ice sheets. b, The mean value of population sizes over the past 450K years. c, The variance of population sizes over the past 450K years. Values are capped at 1500.

We then explored which demographic parameters contributed in generating the population structure observed in leopards. After confirming via power analysis that the data were sufficient to characterise the parameters of interest (Supplementary Fig 2; Supplementary Table 5), we estimated the posterior distributions via ABC-RF (Pudlo et al., 2016; Raynal et al., 2019). Among the seven parameters that define the demography of leopards in our model, we found that migration rate, allometric scaling and allometric factors (parameters linking effective population size to habitat suitability) and initial population size show strong posterior signals, suggesting that they contributed to the differentiation of leopards (Supplementary Fig. 1). The low expansion rate of leopards (maximum a posteriori (MAP) estimate ~ 0.026) indicates that only around 4% of the individuals would expand to neighbouring areas. Low values of allometric scaling parameters (log scaling factor MAP estimate ~ 4.7; scaling exponent MAP estimate ~ 0.37) indicate slow increase and low maximum population sizes per deme for leopards. Such parameters align with our expectation for a territorial, large apex carnivore (Macdonald and Loveridge, 2010). We did not find a strong signal in directed expansion, suggesting random diffusion in this model is sufficient to explain global leopard expansion. The lack of signal in altitude cost, on the other hand, suggests that altitude does not constitute a migration barrier to leopards beyond their effect on local climate, in line with the records of leopards living in mountains (Vernes et al., 2021).

### Establishment of global leopard populations

In our aggregated demography, the initial phase of leopard population dynamics is characterised by a fast expansion out of Africa via the Arabian peninsula, establishing the populations on the entire Asian subcontinent over the course of just tens of thousands of years (Fig. 2). The rapid nature of this expansion led to a strong diversity gradient in Asia through repeated bottlenecks and gene surfing, eventually creating the highly diverse genetic structure observed today. The rapid colonisation process is also in line with previously reported mitochondrial and nuclear population structure, where there is a strong pattern of monophyly across Asian leopards (Paijmans et al., 2021), and potentially European leopards (Paijmans et al., 2018).

The reconstructed demography shows that the west bank to the Red Sea was a corridor for the out-of-Africa dispersal of leopards. Throughout the glacial cycles, a narrow coastal area of the west bank was suitable for leopards, and shows lower population densities (Fig. 2b). During the interglacial periods, the suitable area extended towards the eastern part of the Sahara, potentially allowing for increased gene flow between Africa and Asia. This could explain the pattern of low-level gene flow between Asia and Africa, as suggested by consistent phylogenomic monophyly, and limited evidence for admixture (Paijmans et al. 2021): even during favourable interglacial periods, when the Asian range expanded, the connection between Asia and Africa was narrow; during glacials periods it became minimal.

Whilst we were not able to include the recently sequenced Arabian leopard genome from a captive individual in our model (Mochales-Riano et al., 2023), as its high levels of inbreeding suggest it may not be an accurate reflection of the wild populations, the predicted barriers between Africa and the Middle East are in line with Mochales-Riano et al.’s genetics results showing its high similarity to other Asian genomes.

### Stable climate conditions in Africa led to shallow isolation by distance

To understand the contrast of African and Asian population structure, we further investigated the reconstructed population dynamics by calculating the mean and variation of population sizes for each deme over time (Fig. 3b,c; Supplementary Figure 7, 8). We found that sub-Saharan Africa stood out as having high average and low variance, caused by consistently stable effective population sizes throughout the glacial cycles (Fig. 3a), with the reconstructed population density dropping to near zero only in the equatorial part of Congo and the western part of South Africa at times. This stable inhabitation of the African continent explains the high degree of panmixia and the lack of structure observed in African leopard genomes despite the large geographic distance among them. This result is also in line with previous work by Pecnerova et al (2021) which reported stably high population size in the Pleistocene and Holocene, in addition to the absence of any obvious geographic barrier in Africa.

Morocco is the only area in Africa that shows high levels of fluctuation of population size and reduced connectivity to other parts of Africa. During the LGM (left-bottom figure of Fig. 3a), suitability of the region significantly dropped (Supplementary Figure 8), hinting at possible population bottlenecks and fragmentations during that time. These factors may have collectively contributed to the elevated genetic distance between the Moroccan and other African samples, and potentially also explain the previously reported unexpected phylogenetic pattern in which the Moroccan individual was found to be a sister lineage to all leopards or all Asian leopards in 20% of the genomic windows (Paijmans et al. 2021).

### Asian climate created differentiation patterns

In contrast to Africa, leopard ranges in Asia underwent several contractions and expansions following the glacial cycles, with only south and south-east Asia showing continuous inhabitation (Fig. 3A, 3B, Supplementary Figure 7, 8). In our reconstructions, the much stronger genetic structure observed in Asia, which has led to the recognition of several subspecies, has been shaped by two factors: an initial rapid expansion into the continent, which, via a series of founder effects, created a steep isolation by distance (Fig. 2B) (as discussed in the past for e.g. humans, Ramachandran et al., 2005); and repeated fragmentation of their range during glacial periods (Fig 3).

During glacial periods, northeastern Asia regularly became unsuitable for leopards, with Korea acting as a possible refugium, analogous to what was previously reported for dogs (Kim et al., 2013). Similarly, the species distribution in Afghanistan and central China became fragmented during the glacial periods, especially the LGM. These repeated episodes of fragmentation, based on the isolation by distance pattern resulting from the fast expansion out of Africa, further contributed to the strong genetic differentiation observed among Asian leopards (Fig 1d; Uphyrikina et al., 2001, Paijmans et al., 2021).

Strong isolation also persisted on the eastern bank of the Red Sea, where the southern Arabian Peninsula is connected to Levant through a narrow coastal corridor during glacial periods, but has larger areas of potential connectivity to the Iranian plateau during interglacials phases (Fig 2a). Such strong patterns of repeated isolation could explain the previously reported strong genetic divergence between the Arabian subspecies (*P*.*p. nimr*) and African leopards (Mochales-Riano et al., 2023), despite geographical proximity.

The climate-induced connectivity change between islands and mainland is another possible source contributing to the differentiation of leopard subspecies. A case of this is the Palk Strait, in which the connectivity between mainland and Sri Lanka recurrently emerged in glacial periods, and disappeared during non-glacial phases. Similarly, the connection between Indonesia and mainland Asia was determined by glacial periods, which might have shaped the subspeciation of the Sri Lanka leopard (*P*.*p*.*kotya)*.

We did not model explicitly the genome of the Java leopard, as it is challenging to model the complex connectivity changes in the Indonesian archipelago. However, projecting our model to that area suggested that the island could be reached during periods of lower sea level (Fig 3a), and then became more isolated during interglacials phases; these long periods of isolation might explain the placement of its mitochondrial genome as basal to all extant Asian leopards (Wilting et al., 2016).

### Climate-driven repeated colonisation-extinction events in Europe

Due to the mechanistic nature of the model, it is possible to parameterise the impact of climate on the population dynamics of Africa and Asia, and then project it to Europe, even in the absence of genomic data from European leopards.

In the aggregated demography, Europe was one of the most unstable parts of the leopard range, showing a high population size variance over the past 450,000 years (Fig 3, 4). The fossil record suggests that Europe was first colonised from Asia during the Middle Pleistocene (Turner and Anton, 1997; Baryshnikov, 2011; Kurten, 2017), either via a southern route through Anatolia and the Balkans, or a northern route through the Pontic Steppe (Fig. 4, Diedrich, 2013; Baryshnikov, 2011). Our habitat suitability and demographic reconstructions suggest that the northern route remained disconnected even during the warm interglacial phases, supporting the role of the Anatolian and Balkan peninsulas as the main route for the initial colonisation and any subsequent gene flow of leopards in Europe.

**Figure 4.**
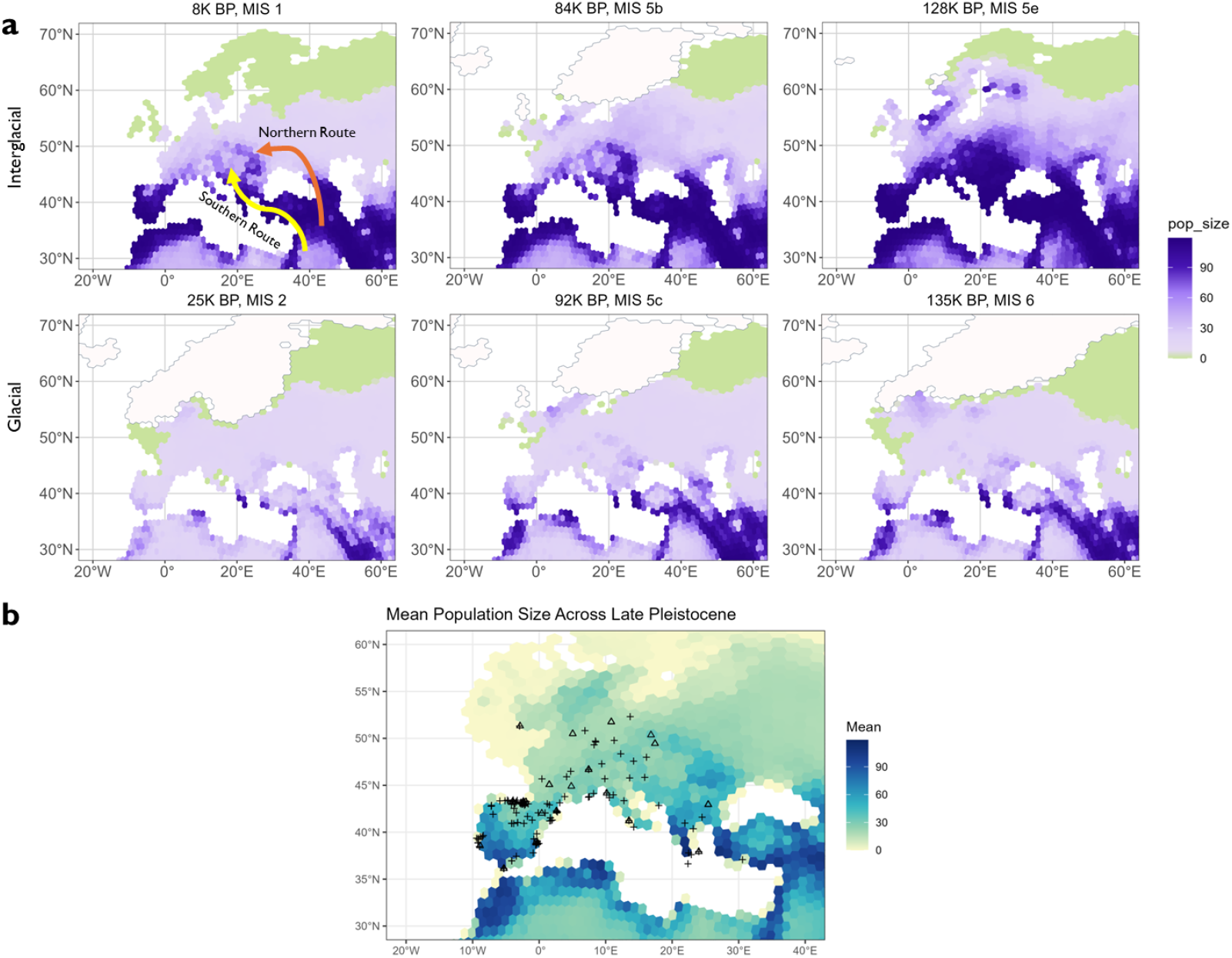
European leopards demography. a. European population size fluctuation during recent glacial cycles. In the first plot, we show the Northern and Southern Route as proposed in Diedrich, 2013. b. Fossil records (crosses) and radiocarbon dates (triangle) records in late Pleistocene. Background is the mean predicted population sizes over the Late Pleistocene (129,000 cal. BP to 11,700 cal. BP).

Our demographic reconstructions show that the majority of Europe was unsuitable during most glacial phases. Interglacial phases, on the contrary, provided brief windows when leopards were able to expand into the subcontinent, before becoming mostly extinct again as climate cooled. The last major window of suitable climate was observed during MIS 5a (90,000-80,000 years BP), when leopards were distributed across large parts of south and south-east Europe. After MIS5 (ca. 128-73 kya), population size in Europe decreased sharply, reaching its lowest point during the LGM (around 21 kya), in line with the extinction of central European leopards around 25 kya based on the fossil record (Diedrich 2013; Sommer & Benecke 2006; Ghezzo & Rook 2015; Nagel 1999; Sauque et al 2016).

To further validate our model in Europe, we compiled a list of published Late Pleistocene European leopard fossils and radiocarbon dated records. We superimposed these past presences on the aggregated demography of the Late Pleistocene (129,000 cal.BP to 11,700 cal. BP) (Fig. 4b; Supplementary Figure 7). Most of the records fell within Southern and Central Europe, where the predicted area has higher average suitability. Notably, the Iberian Peninsula shows the highest density of leopard records, consistent with its higher and longer suitability compared to other parts of Europe, although we caution against overinterpreting this pattern as it is potentially affected by sampling effort.

We further conducted time-series analysis of European leopards using radiocarbon dated records. We compared modelled population size change in demes with dated records against dates density. We found that the number of radiocarbon dates decreased as the LGM approached, coinciding with the population size decrease predicted by our model (Supplementary Figure 9 and 10). This evidence suggests our model reflects possible habitat suitability change in Europe.

Our reconstructions thus suggest that climatic conditions are sufficient to explain the disappearance of European leopards during the LGM. However, our model predicts the potential of leopard resurgence in southern Europe after the LGM, with a peak around ca. 8 ka (Figure 4a), yet no evidence in the fossil record indicates such recolonisation ever occurred. The end of the Pleistocene and the beginning of the Holocene also coincide with increased activity of humans in Europe, which is known to have exerted a large impact on fauna at that time. Humans have been named as potential causes of megafauna extinction for many European species (Bergman et al., 2023; Bartlett et al., 2016; Prescott et al., 2012), thus it seems likely that, whilst climate might have been sufficient to drive the local extinction of the European leopard, anthropogenic impact was the key force that prevented the species return to this continent.

The predicted pattern of repeated expansion and contraction is also consistent with the fossil record before the Late Pleistocene. Diedrich (2013) noted the stark phenotypical differences between leopards or leopards-like *Felidae* in different glacial cycles, some even before the Middle Pleistocene, and hypothesized the existence of multiple leopard subspecies in Europe. The distinctiveness and temporal separation among these subspecies would be consistent with multiple colonisations of Europe, or alternatively a strong bottleneck in a small local refugium during glacial phases, coupled with limited genetic continuity and possible selection under relatively unfavourable conditions. A potential European interglacial refugium was in the southern part of the Iberian Peninsula, where the fossil record also shows a potential persistence of leopard occupation (Sanchis et al., 2015; Sauque et al 2016). However, these hypotheses would require more zooarchaeological and DNA evidence to test.

## Conclusion

Understanding the drivers of past demography dynamics and their evolutionary consequences is a central target in evolutionary studies. In the case of leopards, by combining multiple lines of evidence, namely genetics, ecological surveys of present occurrences, and paleoclimate model outputs, we show that the discordant patterns of leopard genetic structure in Africa and Asia can be explained by the fragmentation of the range in the latter continent during glacial cycles. These results show the importance of generative models to explicitly test the plausibility of scenarios to understand the drivers and barriers of movement.

A further key aspect of our modelling approach is that, despite having no genomic data from Pleistocene European leopards, we were able to investigate the extinction dynamics in that continent. Whilst ancient DNA is required to investigate interglacial recolonisation of Europe from Asia, and the role of a potential Iberian refugium, we were able to show that climate was at least a major driver for the disappearance of European leopards before and during the LGM, but not why leopards did not recolonise Europe afterwards. Thus, by characterising the impact of climate on population dynamics, we can also infer impacts of other crucial factors: in this case, a likely impact of increased human activity in Europe on animal populations.

## Materials and Methods

### An overview of CISGeM

Climate-Informed Spatial Genetics Modelling (CISGeM) is a spatial-temporal framework to simulate demography and corresponding genetics. The model first reconstructs the geographic expansion of species, as well as their spatial density in the unit of effective population size per grid cell. Such simulation processes are controlled by a few demographic parameters determining growth rate, expansion rate, and the link between carrying capacity and suitability inferred by SDM. Genetics simulation in CISGeM is extended from the demographic simulations, during which the genealogy relationship of simulated loci across space and time are recorded. Hence, once a simulation successfully reconstructs all provided empirical samples (that demes with empirical samples have at least 1 individuals at the given time), we can then calculate the expected genetic difference between each pair of simulated haploids (or diploids if empirical samples contain two haploids). By comparing the genetic summary statistics from simulations against the empirical ones from genomic data, we can then evaluate the accuracy of simulations, selecting the ones closer to observed values, and thus obtain demographic history and their drivers from these more reliable simulations.

Compared to demographic inference using only genetic data, the model can recreate scenarios where connectivity between two populations changes due to environmental and geographical factors, allowing the reconstruction of complex meta-population dynamics across time and space. Built on the basis of SDM, the model uses genetics data to validate the reconstructed demography, hence providing robustness over traditional niche modelling.

### Species Distribution Modelling and past suitability reconstructions

In CISGeM, Species Distribution Modelling (SDM) is used to connect leopard suitability with environmental changes in the past, providing the background information simulations needed. We used an approach largely similar to that described in Leedham et al (2023), except for using a different climate dataset (Krapp et al., 2021) to allow for deeper time reconstruction.

We first take the leopard occurrence data used in Leedham et al., 2023, which contains 8,708 occurrences. According to historical leopard habitat, we chose Africa, Eurasia and New Guinea as the background climate, and cropped these areas using R package ‘terra’. To visualise the niche difference between the background and occurrences, we used PCA to project the variables onto a 2D space.

To select the bioclimatic variables for the SDMs, we used the same procedure described in Leedham et al (2023): we first plotted the distribution of all available climate variables of leopard occurrences against that from the background, it is equivalent to the functions plot_pres_vs_bg() and dist_pres_vs_bg() from the R package tidysdm (Leonardi et al. 2024). After this step, we removed highly correlated variables (threshold=0.7) using R package ‘caret’ (version 6.0.94), resulting in 6 variables out of 19 being retained for the SDM: BIO05 (Max temperature of warmest month), BIO07 (Temperature annual range), BIO13 (Precipitation of wettest month), BIO14 (Precipitation of driest month), Leaf Area Index (LAI) and rugosity. PCA was performed to visualise the difference between the niche of selected variables vs the background.

To reduce the sampling density bias, we then thinned the dataset using package spThin (Aiello-Lammens et al., 2015), removing presences that have a distance smaller than 70 km, which leaves 1,361 occurrences records.

We then used the R package Biomod2 (version 4.2-3, Thuiller et al., 2009) to perform the modelling with the following algorithms. We used the landmass of Africa, Eurasia and New Guinea as background from which pseudo-absences were randomly sampled 5 times, with a minimum 150 km radius. We then used R package ‘blockCV’ (version 3.1.2, Valavi et al., 2018) to split the data into 20 longitudinal columns, which were subsequently assigned to one of the five data splits. Models were then run on each of the datasets with cross-validation, selecting each as the test set and the rest four as the training set, using four algorithms: Generalised Linear Model (GLM), Generalised Boosted Model (GBM), Generalised Additive Model (GAM) and Random Forest (RF).

Runs with True Skill Statistic (TSS) over 0.7 were retained and averaged by one of the following methods to build four ensemble models: mean, median, committee average and weighted mean (Araujo and New 2007). After the ensemble, the models were evaluated using get_evaluations().

Once the SDM was built, the model was used with using a palaeoclimatic reconstruction dataset (Krapp et al., 2021) to project the climatic suitability across the past 450,000 years in 1,000 year intervals, using the BIOMOD_Projection() and BIOMOD_EnsembleForecasting() functions. The grid file is projected onto hexagonal grids using cdo -remaplaf command. (Schulzweida, 2023).

### CISGeM Demography Simulation Process

CISGeM uses the suitability estimates from the SDM to condition the demography. Space is represented by a hexagonal grid in this case covering Africa and Eurasia, where local environmental changes according to the SDM. The model uses forward simulations that track the parameter change across time steps to allow local environmental change through time. This changing environment background allows us then to place species dynamics inside the grid, and track the demography change through the changing environment.

The detailed process of demography simulations is given below. CISGeM operates on hexagon grid cells (the distance between the centers of two hexagonal cells is 120.6 ±7.6 km; the variation is due to the earth having a spheroid rather than a perfect spherical shape and the grid not being perfectly regular). Due to sea level change, grids have a binary accessibility that might be varying through time, which is determined by sea level data obtained from climate dataset (Krapp et al., 2021). For simplicity of modelling migrations, we set the grid to be accessible even when only a small proportion of land is present in a grid.

For a given time step of simulation *t* and given grid cell *x*, the carrying capacity *K* (*x,t*) (theoretical maximum number of leopards that can live in a cell) of an accessible cell is given by:

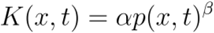

Where *P* (*x,t*) denotes the probability of a species inhabiting cell *x* at tme *t*, and is calculated from the species distribution modelling (see Species Distribution Modelling section). α and β are two parameters: allometric scaling factor and allometric scaling exponent, respectively, that controls the relationship strength between the SDM probability and the carrying capacity.

The simulation is then initiated at selected starting time point *t*_0_ and grid(s) *x*_0_, where the population size *N*(*x*_0,_ *t*_0_) is given by:

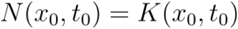

In other words, the initiation is set to the maximum number of individuals at that time and space.

After initiation, CISGeM determines the following population sizes are through two processes: the local growth of populations within the cell, and the spatial migrations from adjacent cells.

Growth model: For cell *x* and time *t*, the population size at next time *N* (*x*, *t* +1) is given by:

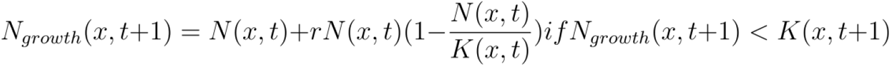

Otherwise, a rescaling strategy is applied to hard cap the number of individuals:

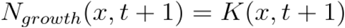

Where *r* is a parameter - intrinsic growth rate.

This growth model hence gives an approximately exponential growth at low population sizes, and saturates when population size is close to the carrying capacity.

Migration model: two types of migration are modelled in CISGeM. The first type migration is a non-directed uniform expansion into neighbour cells, and the second type represents a directed migration driven by resource availability gradient between cells.

For both types of migration, the effect of altitude is considered. Intuitively, a steep altitude change between cells should make migration more difficult. Between cell *x*_1_ and *x*_2_, the altitude scaler is defined as:

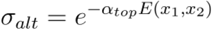

Where α_*top*_ is a parameter, altitude cost, which defines the strength of such altitude change, and *E*(*x*_1,_*x*_*2*_) denotes the absolute altitude change between the center of two cells, as we assume ascend and descend have the same effects.

The first type migration, undirectional migration, is hence defined as following: at time t, from grid cell to *x*_1_ one of its neighbouring cell *x*_2_, the number of individuals migrating *M*_*undirectional*_ is estimated as:

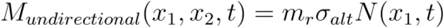

Where *m*_*r*_ is the parameter undirectional migration coefficient. This type of migration is applied to all the neighbouring cells of *x*1.

The second type migration, directional migration, denotes an additional number of individuals moving along the resources gradient. The number of individuals *M*_*undirectional*_ moving from grid cell *x*_1_ to *x*_2_ at time t is hence given by:

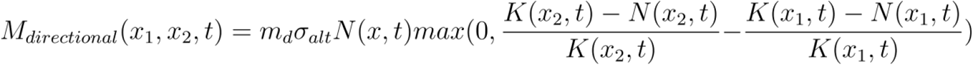

Where *m*_*d*_ denotes the parameter directional migration coefficient, and the term 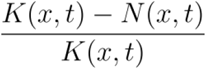 represents the relative availability of resources in the cell x at time t, ranging between 0 (when all the resources are occupied) to 1 (no resources are used). Thus, the number of migrants under this type of migration is proportional to the steepness of the resource gradient.

These two types of migrations can thus be used to examine the possible drivers of migration, that whether patterns can be explained primarily by random motions or complex responses to available sources.

Combining these demographic processes together, the total number of individuals of grid *x* cell at time *t+*1, given its neighbouring cells, can be written as:

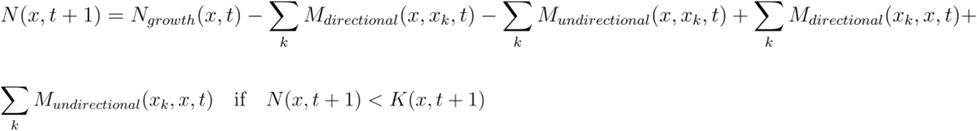

Alternatively, when the final number *N* (*x*, *t* +1) exceeds the carrying capacity *K* (*x*, *t* +1), another rescaling strategy is applied to the final number of individuals to prevent the overflow of individuals in a cell. For this we define that the local growth individuals have higher priority compared to migrants, that is, if adding the migrants cause the overflow of carrying capacity, the excess migrants will be considered ‘dead’ and will not be counted into the total population, representing a local adaptive advantage.

The demography simulation is hence the repeat of the steps defined above until present (0BP). For each time step, the number of individuals in each cell and the number of migrants from both types of migration between grid cells are tracked and eventually used as the output of the demography model.

### CISGeM Genetics

The demography tracks the full genealogy of simulated individuals, which forms the basis for subsequent genetic simulations. This can be broadly seen as a backward Wright-Fisher process: for a simulated sample in cell *x* at time *t*, its ancestral lineage is either randomly assigned to another sample in the same cell *x* at time *t* − 1; or, if the sample is a migrant from neighbouring grid *x*_*k*_, its lineage will be first moved back to *x*_*k*_, then assigned to a random individual there at *t* − 1. As leopard is a diploid species, both haploids are traced in this process. A common ancestor event happens when two lineages are assigned to the same parental gamete, where the two lineages are merged into one lineage. The backward process is repeated until all the lineages are merged have met. If the multiple lineages are not met when the simulation is not initiated, they are instead considered from a single ancestral population with fixed population size *k*_0_, and use a coalescent model to estimate the timing of additional common ancestor events needed to close the tree.

The traced lineage can be seen as the true tree underneath a single locus. Once the tree topology is obtained, mutation is added to the tree using the given mutation rate, following the standard process in *msprime*.

In practice, to reduce the effect of randomness and avoid simulating the recombination process, CISGeM simulates multiple gene trees from the samples, which can be seen as the lineages of independent loci with no linkage disequilibrium, to obtain more realistic genome-wide summary statistics.

### Genetics data processing

The empirical genetic summary statistics required to fit the simulations is prepared as the following. We first downloaded the raw reads of selected leopards (Supp Table 1) from the European Nucleotide Archive and NCBI Nucleotide Database. We applied a similar pipeline as described in Pecnerova et al., 2021. We first trimmed the adapters of raw paired-end reads and merged them using NGMerge, requiring at least 11 base pairs overlapping on termini. We then decided whether to keep unmerged reads based on their reported coverage in the original papers. For low-coverage historical samples (reported average depth < 9), unmerged reads only represent a small portion of the data and often contain higher fractions of contaminants and were thus discarded. Retained merged reads were then mapped using bwa mem (version 0.7.12-r1039) with default parameters to reference genome PanPar1.0 (Kim et al., 2016), with all duplicated reads marked and removed using samtools markdup (ver 1.14). Average mapped coverages were calculated using samtools stats and homemade scripts. For high-coverage data from modern samples, both merged and unmerged reads were kept, separately mapped using bwa-mem to the same reference genome, duplicated removed as above, and merged using samtools merge. After mapping, only reads that mapped to scaffolds with length larger than 1M bp were kept. Software mapDamage (version 2.2.2, Jónsson et al., 2013) were used to estimate the damage level of genomes.

For high-coverage samples, in order to reduce the bias from inaccurate genotypes calling for accurate pairwise π calculation, we further filtered mapped bases, retaining the ones with both base quality and mapping quality >30. We also filtered by mapped depth, removing bases with depth less than 10 or larger than twice of the average depth of the sample. Diploid genotypes were then called using bcftools (version 1.9) mpileup and bcftools call, and then merged using bcftools merge with default parameters.

For low-coverage historical samples and the highly-damaged Afghanistan sample, instead of genotyping diploid genomes, we called the consensus sequence of data to generate a pseudo-haploid genome, using Consensify v0.1.1 (Barlow et al., 2020, https://github.com/jlapaijmans/Consensify). Consensify filters positions for a particular coverage cut-off (at least 3x here) and calls a consensus base based on a user-defined number of matching bases (at least 2 matches here). This pseudo-haplodify strategy reduces genotyping errors in low-coverage data. ANGSD (version 0.940, Korneliussen et al., 2012) was used to extract counts of positions from mapped reads data, which was used as input for Consensify to generate pseudo-haploid genomes. Once the pseudo-haploid genomes were called, we merged these with diploid genomes into vcf format using custom scripts (github link), marking them as ‘0|.’ or ‘1|.’ in vcf files.

### Pairwise π and Isolation-by-distance Calculation

We use genome wide pairwise π between individuals as the summary statistics for the model. To obtain robust whole-genome level π estimation and match the simulations, we first scanned through all scaffolds to get 5,000-bp windows that have at most 15% missing for all samples. We also require at least 100,000 bp distance between windows to avoid linkage disequilibrium. This resulted in 16,900 windows.

For each window, we then calculated between-π between each pair of haplotypes. As samples have different levels of damage, we first calculate the genetic differences using transversion mutations only, and then correct this value with ts/tv ratio 3.3. This ratio was estimated from modern samples in the dataset by first counting the number of transitions and translocation for each sample, obtaining individual ratios, and then taking the average as the final ratio. The resulting π is hence:

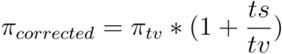

For each pair of haplotypes, we individually consider all overlapping sites to minimise the effect of coverage difference between genomes. We compare each pair of haploids, meaning that when we compare pseudo-haploids with diploid genomes, we use the average of the 2 comparisons as the estimated pairwise π; for diploid samples compared with diploid samples, we take the average of 4 possible comparisons between haploids as the π between two samples. Once the window-level pairwise π are calculated, we used the mean of all π across all windows as empirical π between samples.

Isolation-by-distance was calculated using the mean pairwise π calculated above. The geographical distance between samples was calculated using R package ‘geoGraph’ (version 1.1-1, Jombart et al., 2023). Here, samples were put in approximately 40,000 grids of the earth surface, and the distance between samples are the total lengths of the shortest paths through land grids between each pair of samples. Standard major axis (SMA) regression with 1,000 permutations for each continent were performed with R package ‘lmodel2’ (version 1.7-3, Legendre 2018) to retrieve the strength and significance of the relationship.

### Spatial-temporal simulation with climate information

We simulated a model of out-of-Africa expansion beginning from 450,000 years BP, taking an average of previous estimates in Paijmans et al., 2021 and Pecnerova et al., 2021. Leopard samples with geographical information available were assigned to their recorded locations, and for the samples without explicit geographical information, we assigned them to the middle of the range of their subspecies. We set the generation date of leopards to 5 years. Mutation rate for the Model was set as 1.1e-9 following previous practice in Paijmans et al., 2021. We selected central-east Africa as the starting deme following previous hypothesis (Paijmans et al., 2021), and used a Monte Carlo Interval to sample the following demographic parameters: initial deme radius from 5 to 10 hexagons, directed expansion coefficient [0, 0.014], undirected expansion coefficient [0,0.05], allometric scaling factor [20,500], allometric_sclaeing_exponent [0.1,1], and altitude cost [0,0.3].

The selection of starting population size is crucial for determining the initial population diversity level before expansion. To reduce the parameter range, for each model we first run a small set of simulations to fit the initial size from coalescence, then use abcrf (see next section) to determine the posterior range, and then set the Monte Carlo Interval around the posterior range. We eventually set the interval as [75000,130000].

We performed 90, 224 simulations, of which 51,539 simulations have produced sufficient results that allow at least 1 individual present in the grids that contain empirical samples.

### Approximate Bayesian Computation and goodness-of-fit

To assess whether the simulation captured the desired genetics pattern, we first manually checked that each empirical summary statistics is covered in the range of simulations. With that being satisfied, to better check that global population structure is also captured by simulations, we used gfit-pca from R package ‘abc’ (version 2.2.1, Csillery et al., 2012) to perform dimension reduction on all the simulations generated by individual models, then projected the empirical summary statistics onto the space constructed.

We used R package abcrf (ver 1.8.1, Pudlo et al., 2016) to perform ABC and estimate the posterior probability of sampled parameters. For each parameter, we train a forest of 1,000 trees that fit all pairwise π.

To validate the effectiveness of the models, we performed power analysis by taking out the first 1,000 simulations as a test set, and then used the remaining simulations to train predictive forests for each of the chosen parameters. We then compared the predicted mean and median values, and calculated the R2, the median of the standardised error, the square root of standardised error, and the coverages.

### Analysing Representative Demography

The output of models are presented in spatial-temporal discrete data of demography (the population size per grid), migration (number of migrants between grids) and common ancestors (tracing the number of lineages). To select the best demography from all simulations, we use abc (R package abc, version 2.2.1, Csillery et al., 2012) to calculate the Euclidean distance between summary statistics of each simulation and empirical values. We then selected the best 1,000 simulations that have lowest distances and extracted spatial-temporal demography information of these simulations. Due to the limitation of computation, we selected demography every 1,000 years. The mean of these best demography is then calculated to represent the reconstructed demography that matches the observed genetics.

To assess the stability of suitability in different regions, we calculated the mean and variance of the effective size of leopards across all reconstructed time periods using the mean of the best simulated demography.

The MIS period data used in Fig.3 is obtained from R package pastclim (Leonardi et al., 2023). For the delta18O we used Lisiecki and Raymo 2005 LR04 Benthic Stack data, downloaded from https://lorraine-lisiecki.com/LR04stack.txt.

### European Late Pleistocene Leopards fossil and date records

We collected published fossil and radiocarbon dates associated with leopard presence in Europe between 70,000 (OR MAXIMUM AGE) and 10,000 (OR MINIMUM AGE) years ago (table). The dates were direct (i.e. obtained from leopard remains) and indirect (associated with archaeological layers in which leopard remains have been found). These have been calibrated with the Bchron R package (Haslett et al. 2008) using the IntCal20 curve (Reimer et al. 2020).

### Plotting

All visualisations were performed with R (version 4.1.2). In most cases, R package ggplot2 (version 3.5.1, Wickham, 2011) was used to create the plots. The land polygon in Figure 1.A used land polygon data from naturalearth database, downloaded with rnatualearth package. All spatial data were stored as ‘sf’ objects (Pebesma, 2018) and plotted using geom_sf() function from R package ggplot2.

## Supporting information

Supplementary Media 1

## Code Availability

The code used for generating input data, running simulations, analysis and plotting are available at https://github.com/EvolEcolGroup/Leopards_project. CISGeM is available at https://github.com/EvolEcolGroup/cisgem. A chaperon package for analysing CISGeM output data is available at https://github.com/EvolEcolGroup/rcisgem.

## Acknowledgements

Z.X was funded by CSC-Cambridge Trust Scholarship. J.L.A.P was funded by the Marie Skłodowska-Curie individual fellowship “RESOURCEFUL” (101028348). A.V.P. is supported by the Natural Environment Research Council grant number: NE/S007164/1. A.M. and M.L. was funded by the ERC Consolidator Grant 647787 ‘LocalAdaptation’ and Leverhulme Research Grant RPG-2020-317. M.C. was funded by the Lise Meitner Pan-African Evolution Research Group. C.P.I. is funded by Emmanuel College, University of Cambridge. A. Hovhannisyan acknowledges funding by EU MSCA-IF under grant agreement 101063265.

## References

Aiello-Lammens, M.E., Boria, R.A., Radosavljevic, A., Vilela, B. and Anderson, R.P., 2015. spThin: an R package for spatial thinning of species occurrence records for use in ecological niche models. Ecography, 38(5), pp.541–545.

Barlow, A., Hartmann, S., Gonzalez, J., Hofreiter, M., & Paijmans, J. L. (2020). Consensify: a method for generating pseudohaploid genome sequences from palaeogenomic datasets with reduced error rates. Genes, 11(1), 50.

Bartlett, L. J., Williams, D. R., Prescott, G. W., Balmford, A., Green, R. E., Eriksson, A., … & Manica, A. (2016). Robustness despite uncertainty: regional climate data reveal the dominant role of humans in explaining global extinctions of Late Quaternary megafauna. Ecography, 39(2), 152–161.

Baryshnikov, G. F. (2011). Pleistocene Felidae (Mammalia, Carnivora) from the Kudaro paleolithic cave sites in the Caucasus. In Proceedings of the Zoological Institute Russian Academy of Science (Vol. 315, No. 3, pp. 197–226).

Bergman, J., Pedersen, R. Ø., Lundgren, E. J., Lemoine, R. T., Monsarrat, S., Pearce, E. A., … & Svenning, J. C. (2023). Worldwide Late Pleistocene and Early Holocene population declines in extant megafauna are associated with Homo sapiens expansion rather than climate change. Nature Communications, 14(1), 7679.

Csilléry, K., François, O., & Blum, M. G. (2012). abc: an R package for approximate Bayesian computation (ABC). Methods in ecology and evolution, 3(3), 475–479.

Danecek, P., Bonfield, J.K., Liddle, J., Marshall, J., Ohan, V., Pollard, M.O., Whitwham, A., Keane, T., McCarthy, S.A., Davies, R.M. and Li, H., 2021. Twelve years of SAMtools and BCFtools. Gigascience, 10(2), p.giab008.

Delser, P. M., Krapp, M., Beyer, R., Jones, E. R., Miller, E. F., Hovhannisyan, A., … & Manica, A. (2021). Climate and mountains shaped human ancestral genetic lineages. BioRxiv, 2021–07.

Ghezzo, E. and Rook, L., 2015. The remarkable Panthera pardus (Felidae, Mammalia) record from Equi (Massa, Italy): taphonomy, morphology, and paleoecology. Quaternary Science Reviews, 110, pp.131–151.

Haslett, J., & Parnell, A.C. (2008). A simple monotone process with application to radiocarbon-dated depth chronologies. Journal of the Royal Statistical Society: Series C (Applied Statistics), 57(4), 399–418.

Jensen, A. J., Cove, M. V., Goldstein, B. R., Kays, R., McShea, W., Pacifici, K., … & Kierepka, E. (2024). Geographic barriers but not life history traits shape the phylogeography of North American mammals. Global Ecology and Biogeography, e13875.

Jónsson, H., Ginolhac, A., Schubert, M., Johnson, P. L., & Orlando, L. (2013). mapDamage2. 0: fast approximate Bayesian estimates of ancient DNA damage parameters. Bioinformatics, 29(13), 1682–1684.

Kim, S. I., Park, S. K., Lee, H., Oshida, T., Kimura, J., Kim, Y. J., … & Min, M. S. (2013). Phylogeography of K orean raccoon dogs: implications of peripheral isolation of a forest mammal in E ast A sia. Journal of Zoology, 290(3), 225–235.

Kim, S., Cho, Y. S., Kim, H. M., Chung, O., Kim, H., Jho, S., … & Yeo, J. H. (2016). Comparison of carnivore, omnivore, and herbivore mammalian genomes with a new leopard assembly. Genome biology, 17, 1–12.

Kitchener, A. C., Breitenmoser-Würsten, C., Eizirik, E., Gentry, A., Werdelin, L., Wilting, A., … & Tobe, S. (2017). A revised taxonomy of the Felidae: The final report of the Cat Classification Task Force of the IUCN Cat Specialist Group. Cat News. Available at https://repository.si.edu/bitstream/handle/10088/32616/A_revised_Felidae_Taxonomy_CatNews.pdf.

Korneliussen, T.S., Albrechtsen, A. and Nielsen, R., 2014. ANGSD: analysis of next generation sequencing data. BMC bioinformatics, 15(1), pp.1–13.

Kurten, B. (2017). Pleistocene mammals of Europe. Routledge.

Leedham, S., Paijmans, J. L., Manica, A., & Leonardi, M. (2023). Niche conservatism in a generalist felid: low differentiation of the climatic niche among subspecies of the leopard (Panthera pardus). bioRxiv, 2023-01.

Leonardi, M., Hallett, E. Y., Beyer, R., Krapp, M., & Manica, A. (2023). pastclim 1.2: an R package to easily access and use paleoclimatic reconstructions. Ecography, 2023(3), e06481.

Li, H., Handsaker, B., Wysoker, A., Fennell, T., Ruan, J., Homer, N., Marth, G., Abecasis, G., Durbin, R. and 1000 Genome Project Data Processing Subgroup, 2009. The sequence alignment/map format and SAMtools. bioinformatics, 25(16), pp.2078–2079.

Li, H. and Durbin, R., 2009. Fast and accurate short read alignment with Burrows–Wheeler transform. bioinformatics, 25(14), pp.1754–1760.

Li, H. and Durbin, R., 2010. Fast and accurate long-read alignment with Burrows–Wheeler transform. Bioinformatics, 26(5), pp.589–595.

Macdonald, D.W. and Loveridge, A.J. eds., 2010. The biology and conservation of wild felids (Vol. 2). Oxford University Press.

Miller, E. F., Leonardi, M., Xue, Z., Beyer, R., Krapp, M., Somveille, M., … & Manica, A. (2021). Post-glacial expansion dynamics, not refugial isolation, shaped the genetic structure of a migratory bird, the yellow warbler. BioRxiv, 2021–05.

Mochales-Riaño, G., Fontsere, C., de Manuel, M., Talavera, A., Burriel-Carranza, B., Tejero-Cicuéndez, H., … & Carranza, S. (2023). Genomics reveals introgression and purging of deleterious mutations in the Arabian leopard (Panthera pardus nimr). Iscience, 26(9).

Padilla-Iglesias, C., Xue, Z., Leonardi, M., Paijmans, J. L., Colucci, M., Hovhannisyan, A., … & Manica, A. (2025). Pan-African metapopulation model explains Homo sapiens genetic and morphological evolution. bioRxiv, 2025–05.

Paijmans, J.L., Barlow, A., Förster, D.W., Henneberger, K., Meyer, M., Nickel, B., Nagel, D., Worsøe Havmøller, R., Baryshnikov, G.F., Joger, U. and Rosendahl, W., 2018. Historical biogeography of the leopard (Panthera pardus) and its extinct Eurasian populations. BMC Evolutionary Biology, 18(1), pp.1–12.

Paijmans, J.L., Barlow, A., Becker, M.S., Cahill, J.A., Fickel, J., Förster, D.W., Gries, K., Hartmann, S., Havmøller, R.W., Henneberger, K. and Kern, C., 2021. African and Asian leopards are highly differentiated at the genomic level. Current Biology, 31(9), pp.1872–1882.

Pebesma, E. J. (2018). Simple features for R: standardized support for spatial vector data. R J., 10(1), 439.

Pečnerová, P., Garcia-Erill, G., Liu, X., Nursyifa, C., Waples, R.K., Santander, C.G., Quinn, L., Frandsen, P., Meisner, J., Stæger, F.F. and Rasmussen, M.S., 2021. High genetic diversity and low differentiation reflect the ecological versatility of the African leopard. Current Biology, 31(9), pp.1862–1871.

Pilowsky, J. A., Brown, S. C., Llamas, B., Van Loenen, A. L., Kowalczyk, R., Hofman-Kamińska, E., … & Fordham, D. A. (2023). Millennial processes of population decline, range contraction and near extinction of the European bison. Proceedings of the Royal Society B, 290(2013), 20231095.

Pudlo, P., Marin, J. M., Estoup, A., Cornuet, J. M., Gautier, M., & Robert, C. P. (2016). Reliable ABC model choice via random forests. Bioinformatics, 32(6), 859–866.

Ramachandran, S., Deshpande, O., Roseman, C. C., Rosenberg, N. A., Feldman, M. W., & Cavalli-Sforza, L. L. (2005). Support from the relationship of genetic and geographic distance in human populations for a serial founder effect originating in Africa. Proceedings of the National Academy of Sciences, 102(44), 15942–15947.

Raynal, L., Marin, J. M., Pudlo, P., Ribatet, M., Robert, C. P., & Estoup, A. (2019). ABC random forests for Bayesian parameter inference. Bioinformatics, 35(10), 1720–1728.

Reimer PJ, Austin WEN, Bard E, et al. The IntCal20 Northern Hemisphere Radiocarbon Age Calibration Curve (0–55 cal kBP). Radiocarbon. 2020;62(4):725–757. doi:10.1017/RDC.2020.41

Sanchis, A., Tormo, C., Sauqué, V., Sanchis, V., Díaz, R., Ribera, A. and Villaverde, V., 2015. Pleistocene leopards in the Iberian Peninsula: New evidence from palaeontological and archaeological contexts in the Mediterranean region. Quaternary Science Reviews, 124, pp.175-208.\

Schulzweida, Uwe. (2023). CDO User Guide (2.3.0). Zenodo. 10.5281/zenodo.10020800

Stein, A. B., & Hayssen, V. (2013). Panthera pardus (Carnivora: Felidae). Mammalian Species, 45(900), 30–48.

Thuiller, W., Lafourcade, B., Engler, R. and Araújo, M.B., 2009. BIOMOD–a platform for ensemble forecasting of species distributions. Ecography, 32(3), pp.369–373.

Turner A, Antón M. 1997. The big cats and their fossil relatives: an illustrated guide to their evolution and natural history. Columbia University Press.

Uphyrkina, O., Johnson, W. E., Quigley, H., Miquelle, D., Marker, L., Bush, M., & O’Brien, S. J. (2001). Phylogenetics, genome diversity and origin of modern leopard, Panthera pardus. Molecular ecology, 10(11), 2617–2633.

Valavi, R., Elith, J., Lahoz-Monfort, J.J. and Guillera-Arroita, G., 2018. blockCV: An r package for generating spatially or environmentally separated folds for k-fold cross-validation of species distribution models. Biorxiv, p.357798.

Vernes, K., Rajaratnam, R., & Dorji, S. (2022). Patterns of species co-occurrence in a diverse Eastern Himalayan montane carnivore community. Mammal Research, 67(2), 139–149.

Werdelin, L., Yamaguchi, N., Johnson, W. E., & O’Brien, S. J. (2010). Phylogeny and evolution of cats (Felidae). Biology and conservation of wild felids, 59–82.

Wickham, H. (2011). ggplot2. Wiley interdisciplinary reviews: computational statistics, 3(2), 180–185.

Wilting, A., Patel, R., Pfestorf, H., Kern, C., Sultan, K., Ario, A., Peñaloza, F., Kramer-Schadt, S., Radchuk, V., Foerster, D.W. and Fickel, J., 2016. Evolutionary history and conservation significance of the Javan leopard Panthera pardus melas. Journal of Zoology, 299(4), pp.239–250.

