## Supplementary figures and images for "Climate Shaped the Global Population Structure of Leopards and their Extinction in Europe"

### Supplementary Media 1

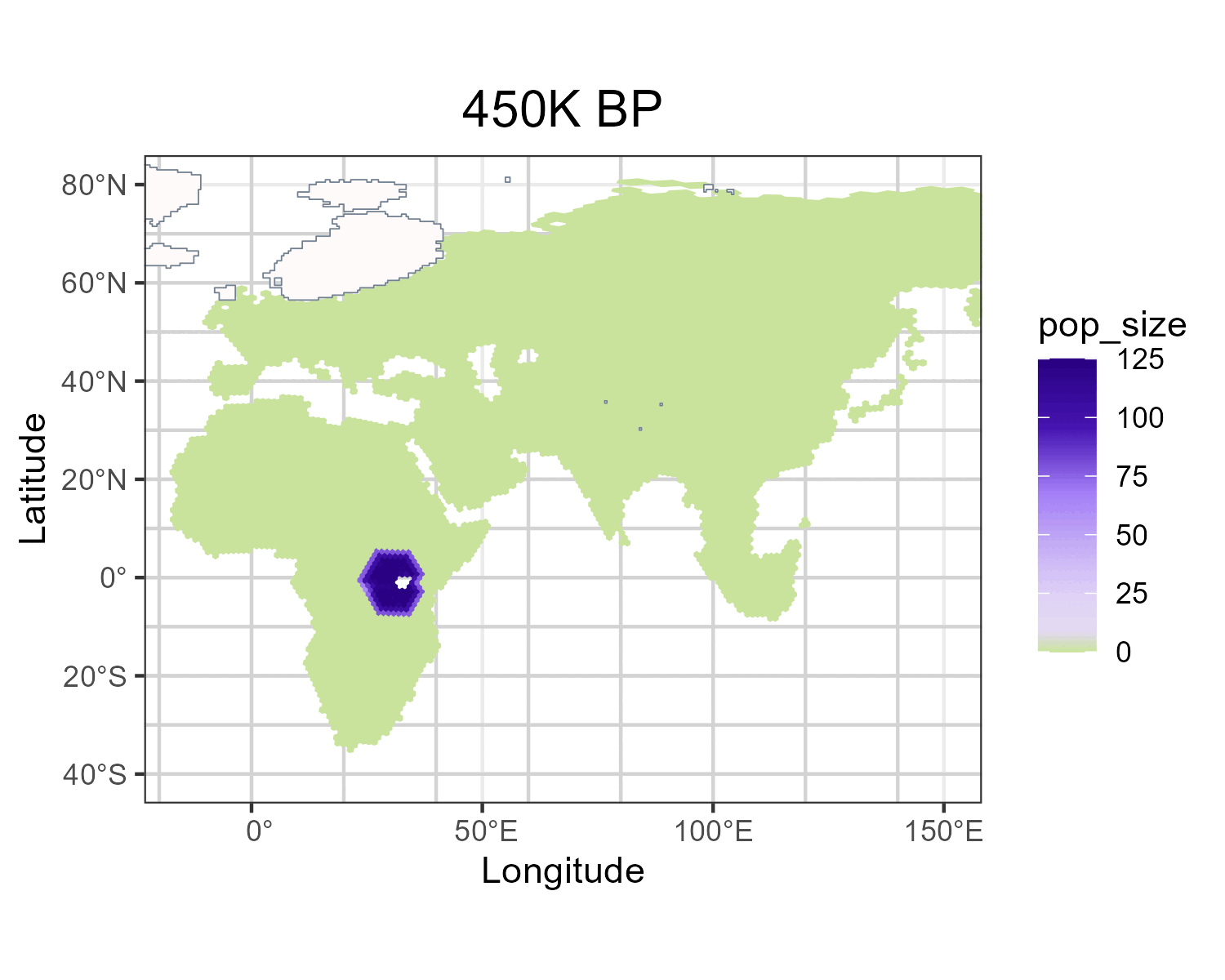
