## Supplementary File 1 for "Climate Shaped the Global Population Structure of Leopards and their Extinction in Europe"

**Supplementary Figures**

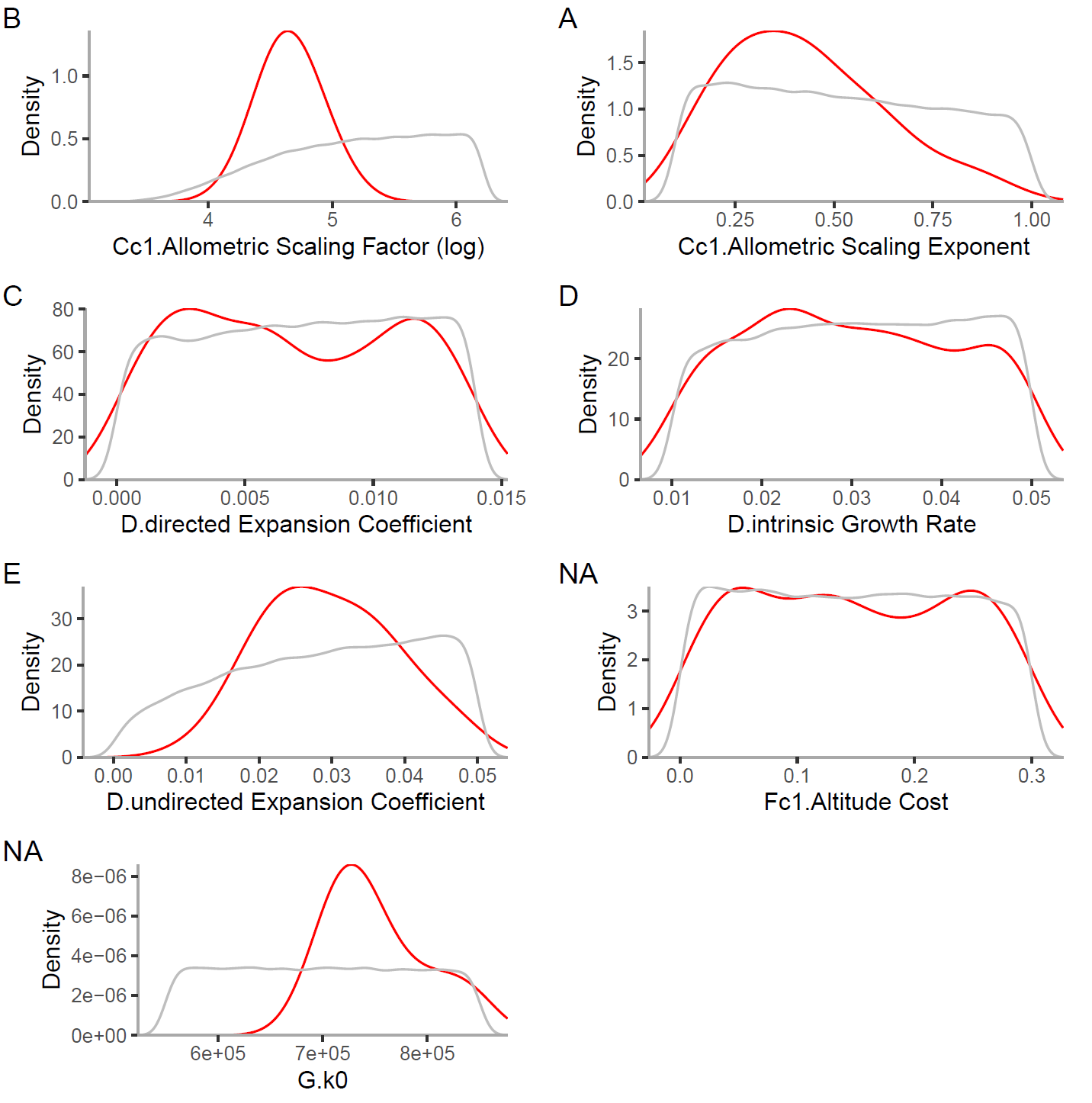

**Supplementary Figure 1.** The Prior (shown in grey) and Posterior (shown in red) curves for the parameters used in ABC.

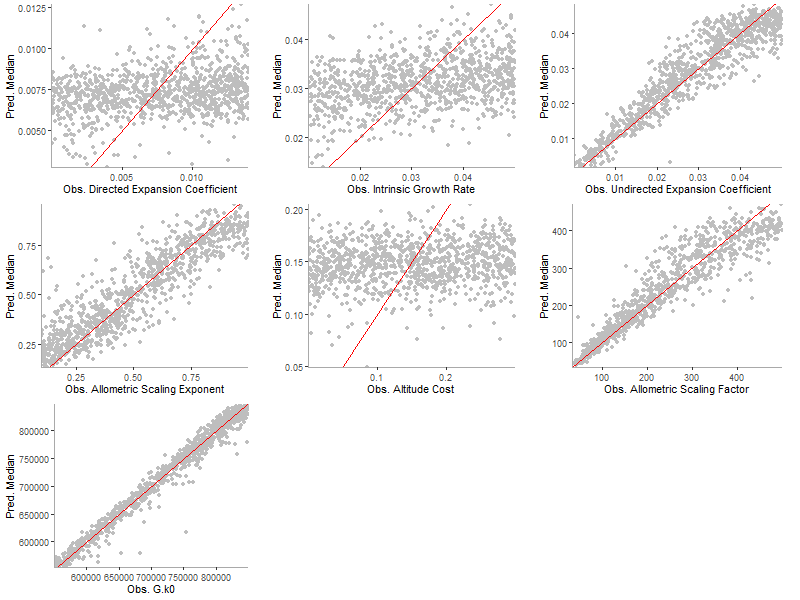

**Supplementary Figure 2.** The power analysis of parameters used in simulations. On X-axes empirical values of parameters used in simulations are shown, and predicted values are shown on Y-axes.

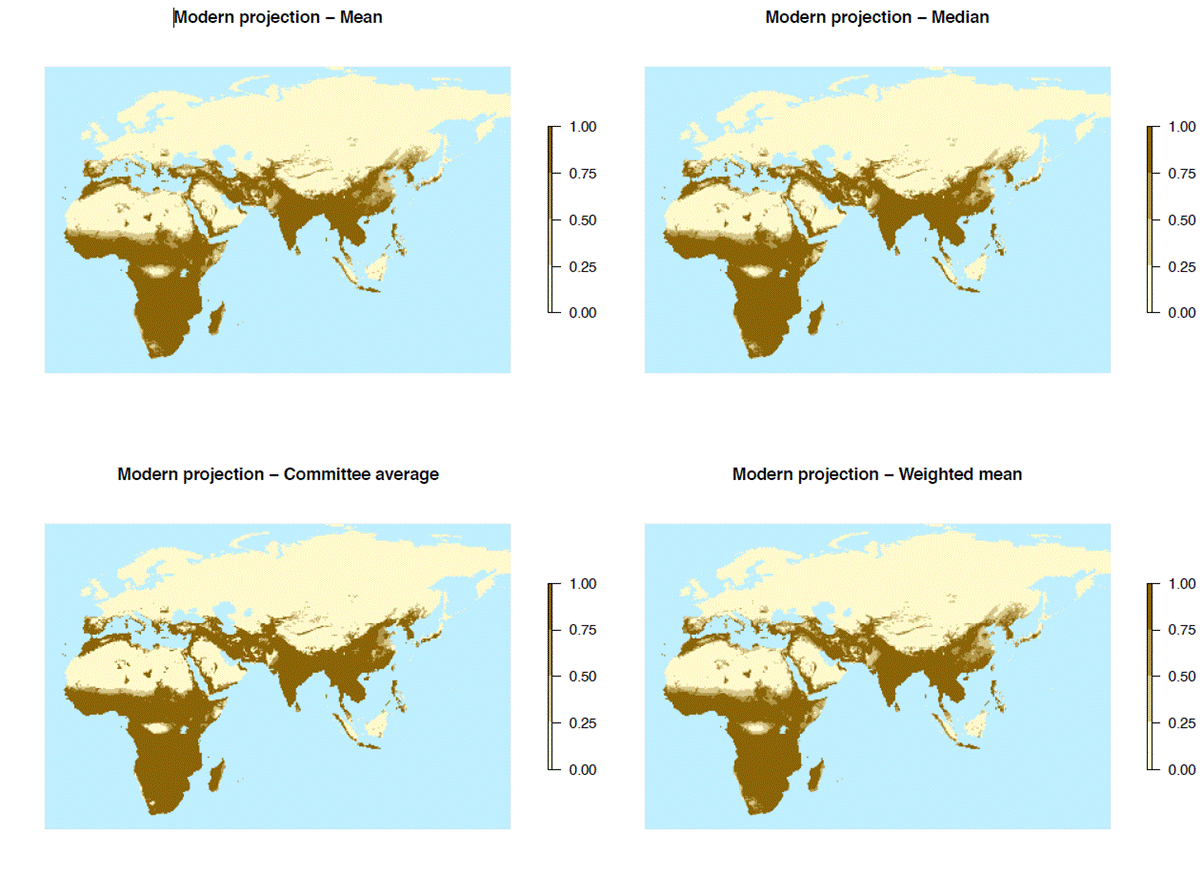

**Supplementary 3.** The projections produced by SDM ensemble models, averaged using mean, median, committee average and weighted mean.

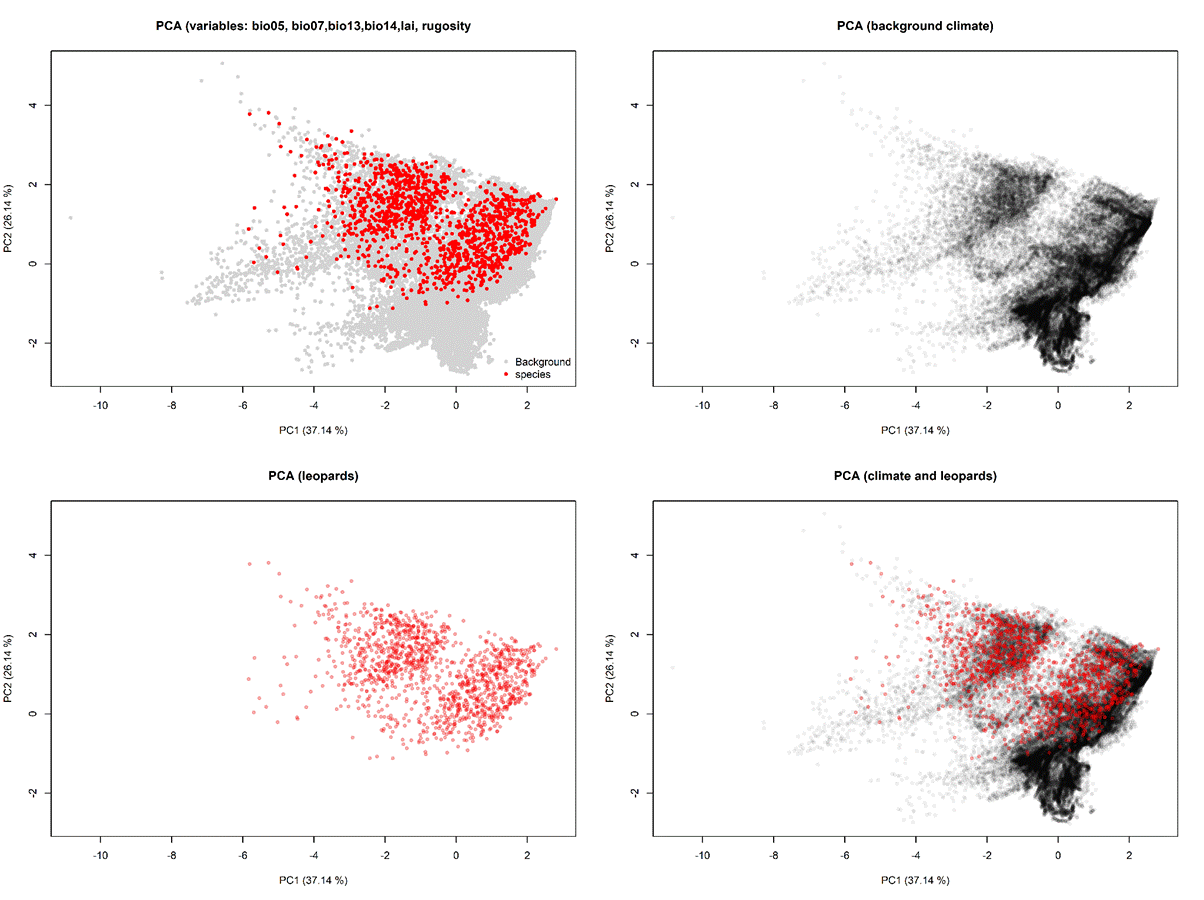

**Supplementary 4.** PCA of the selected climate space of background and occurrences.

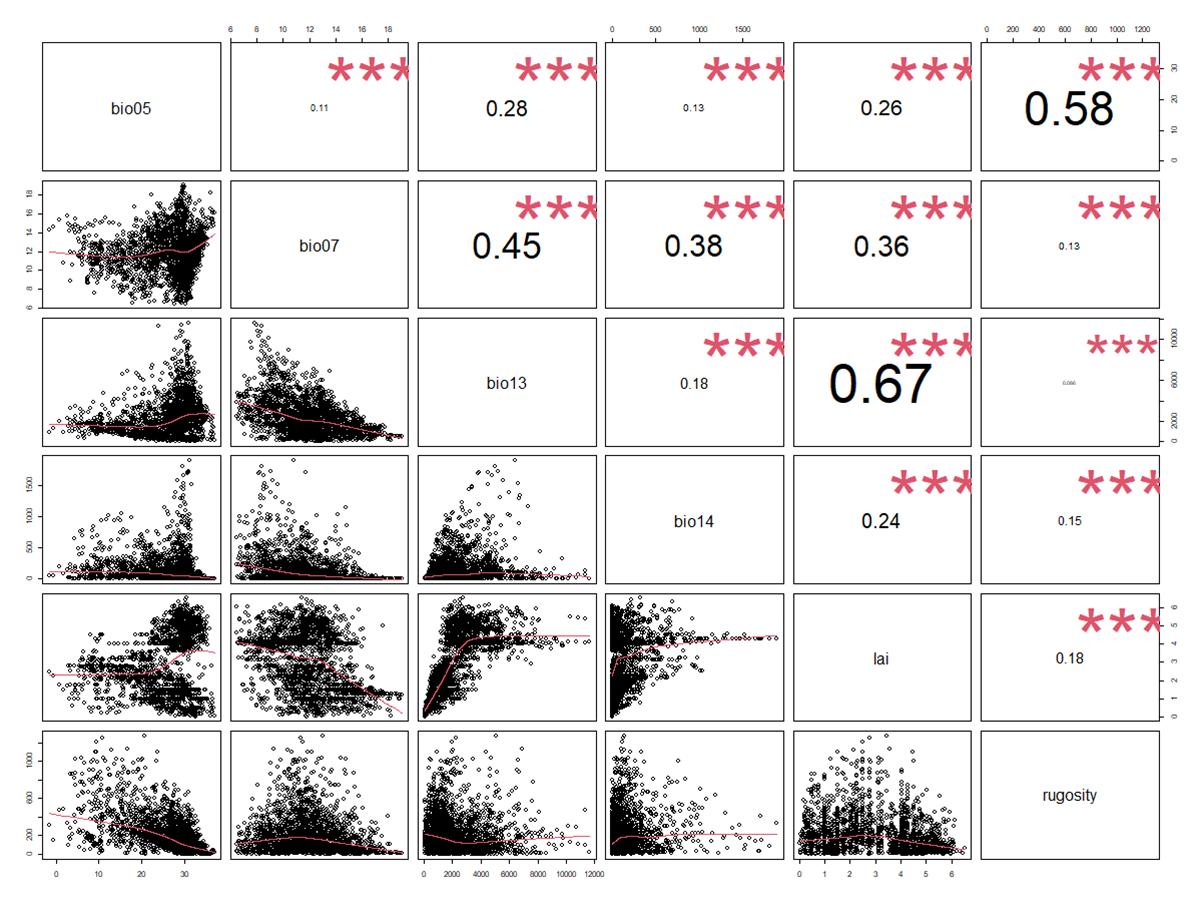

**Supplementary Figure 5.** The correlation between selected variables for SDM.

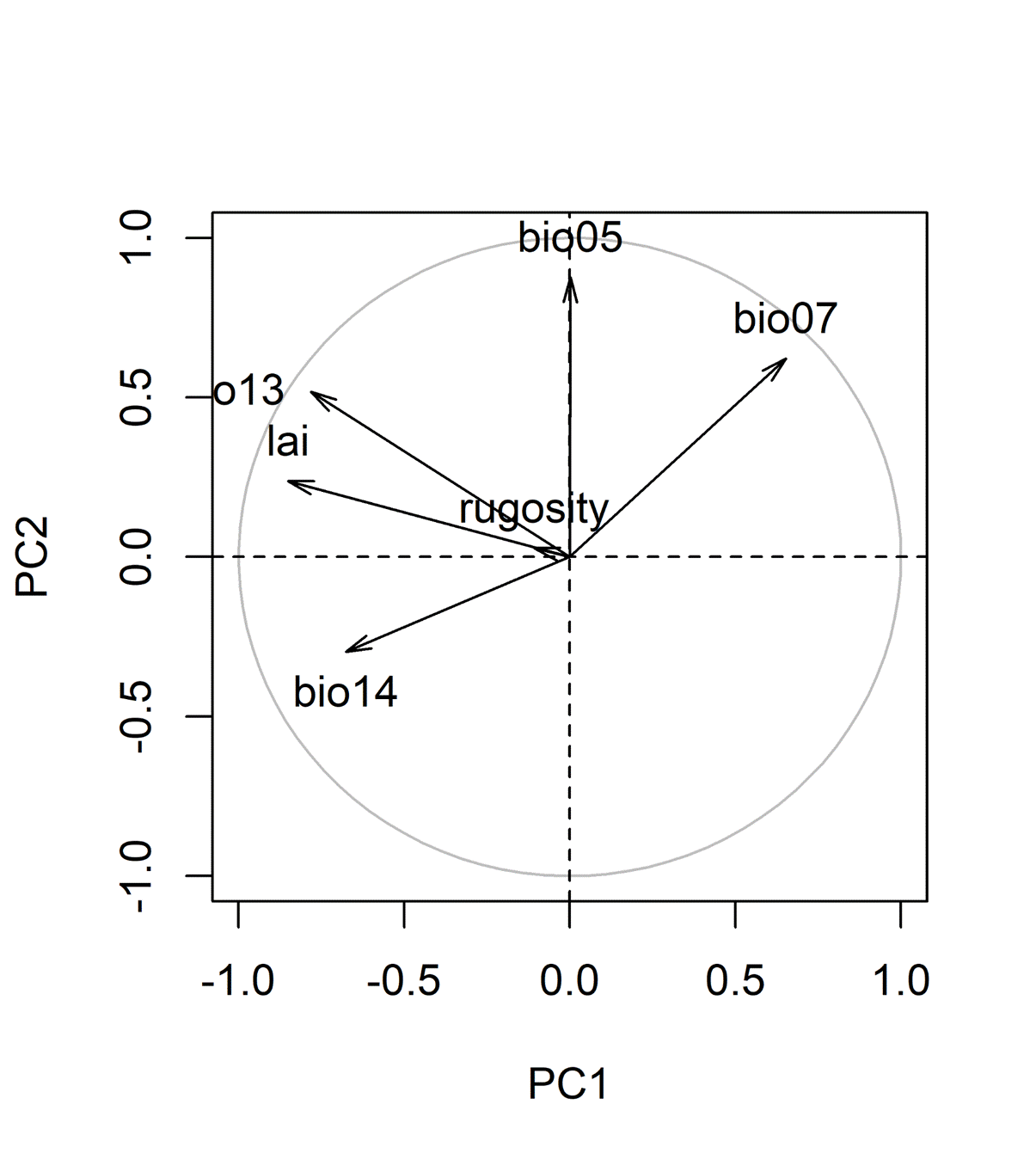

**Supplementary Figure 6.** The directions of selected variables on first 2 PCs.

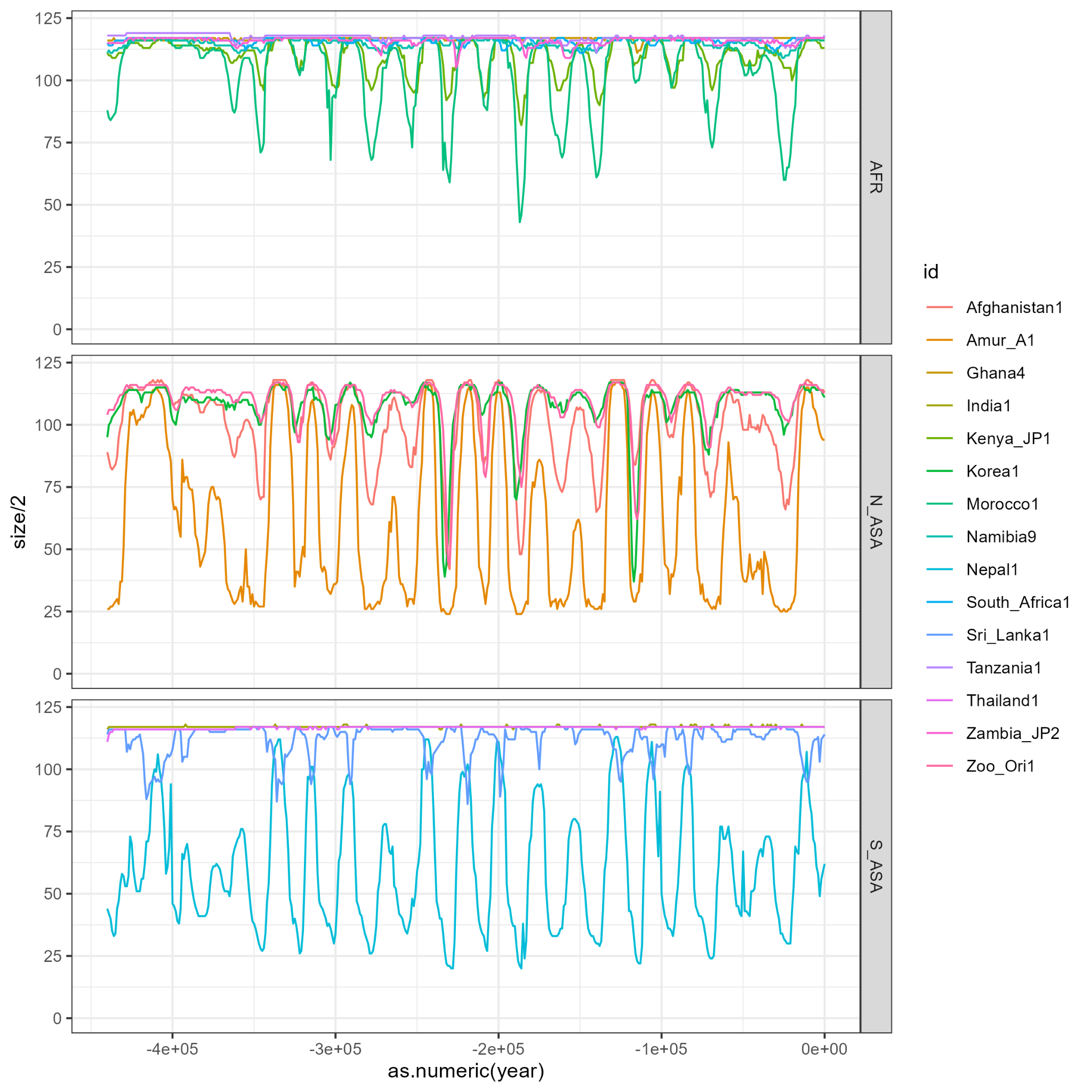

**Supplementary Figure 7.** Mean population size change over the best 1,000 simulations in the cells with empirical samples over the past 450,000 years.

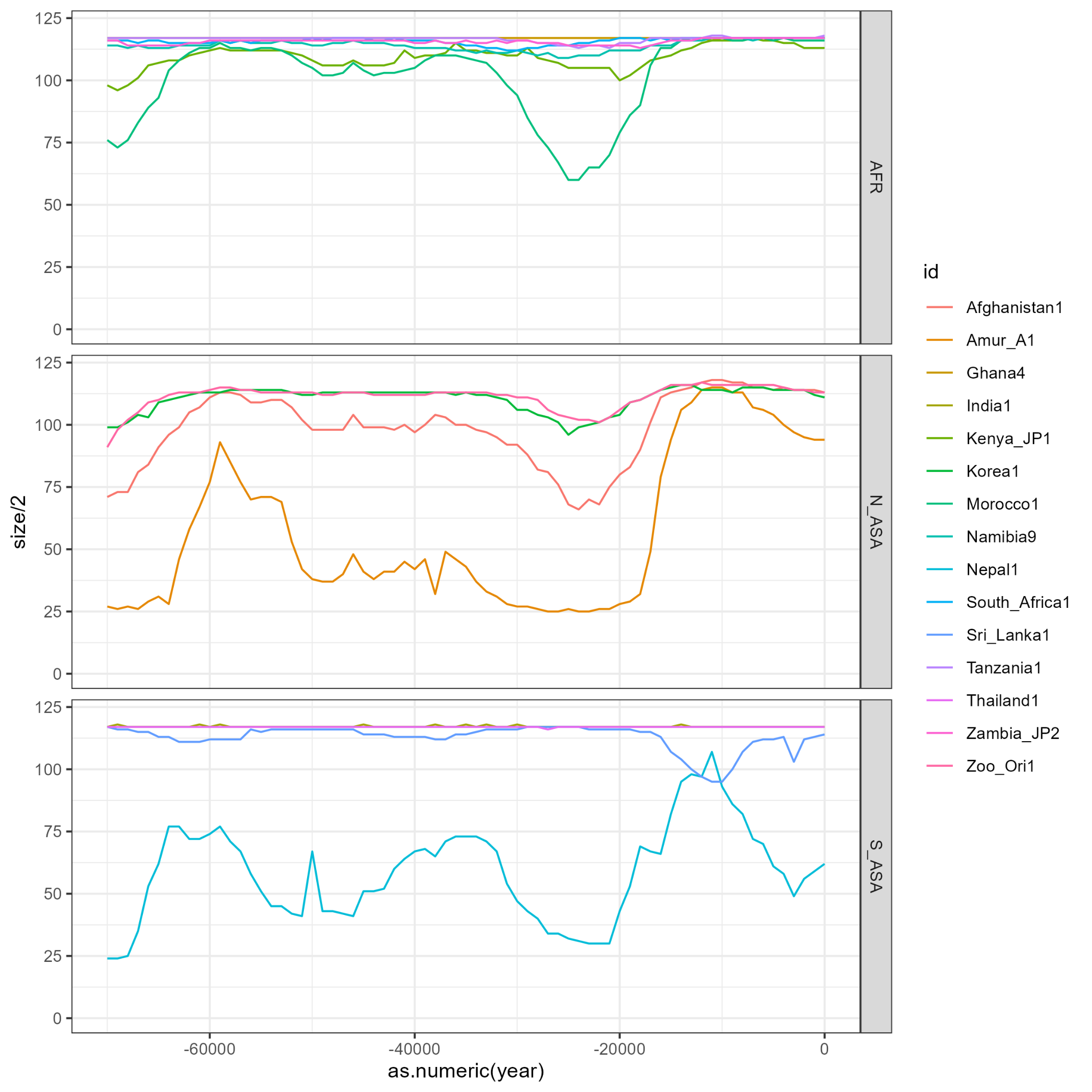

**Supplementary Figure 8.** The change of mean population size of the best 1,000 simulations in the cells containing empirical samples since late Pleistocene (70kya~today).

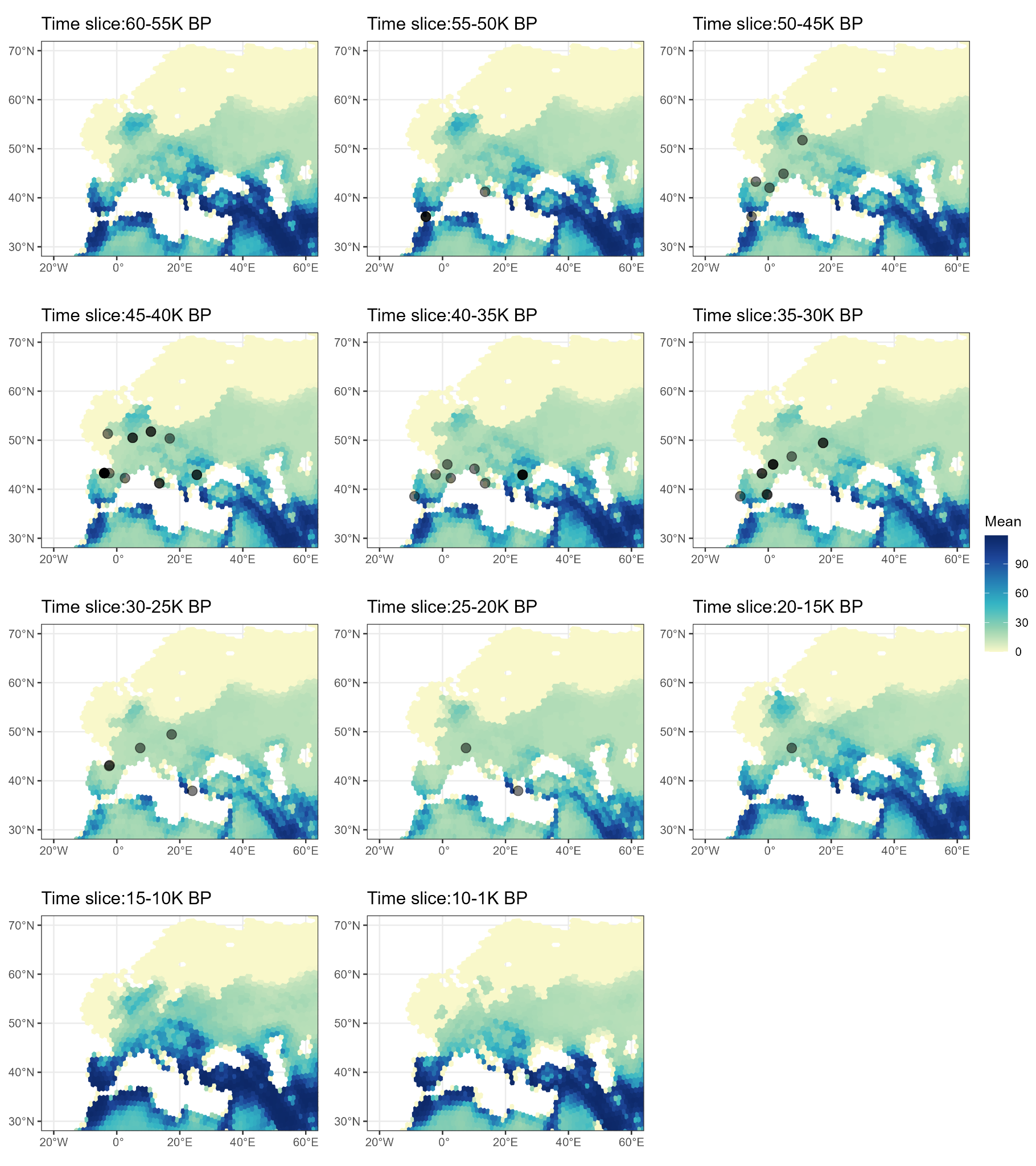

**Supplementary Figure 9.** Spatial and temporal distribution of radiocarbon dates of European leopards. Black dots indicate median calibrated dates in selected time slices. Background colour denotes the mean demography size during the chosen time range.

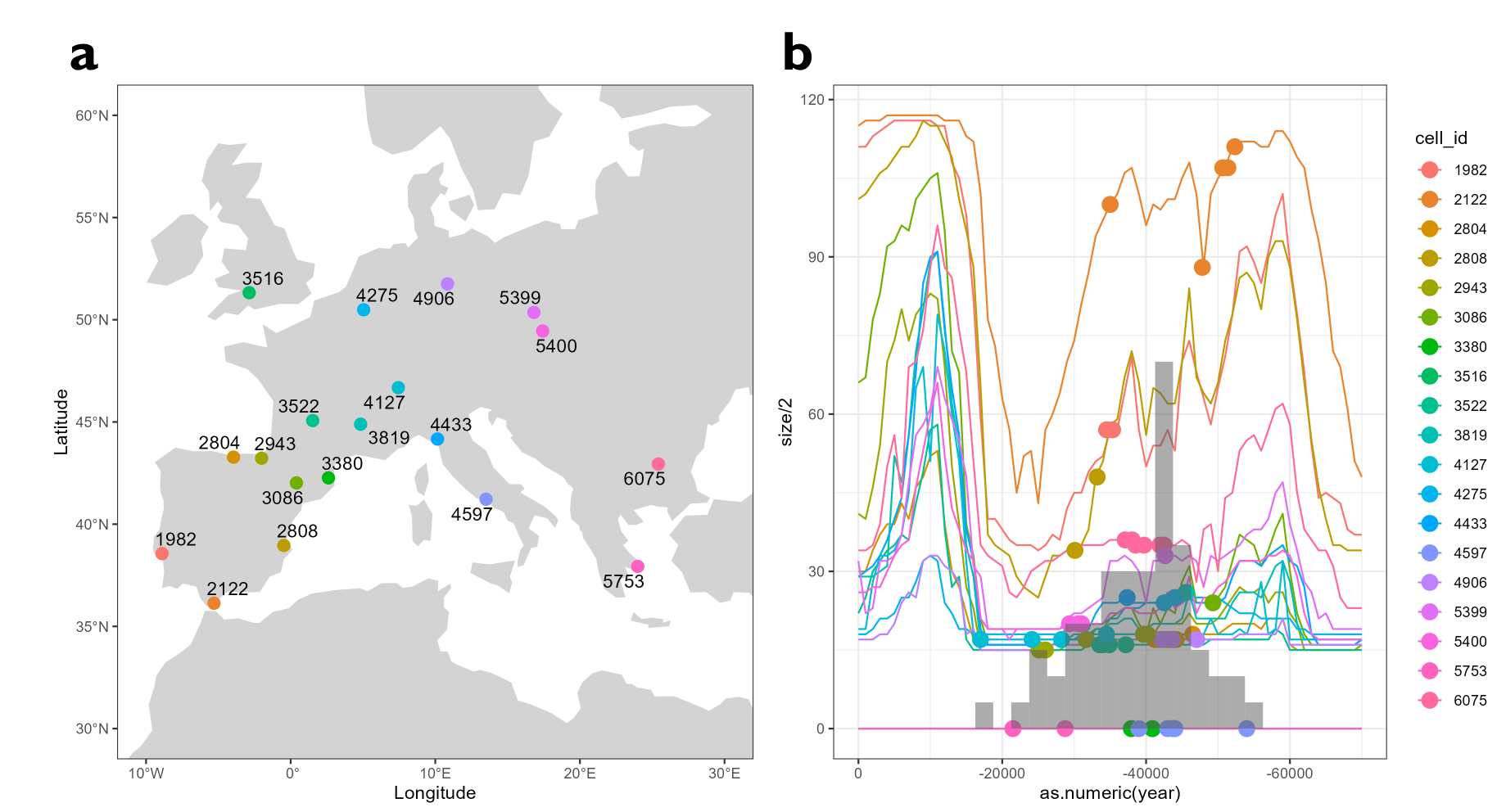

**Supplementary Figure 10.** a. The cells (numbers indicate their ids) that nearest to the location of radiocarbon-dated leopards’ samples. b. Population size changes for each cell in a. Each line denotes the average size change among the best 1,000 simulations during last 70,000 years. Coloured points on the curve indicate median calibrated dates of samples nearest to that cell. Background histogram indicates the density of median calibrated dates in 2,500 years bins. For visualization, density value is timed by 5.

**Supplementary Media 1.** (See online supplementary files) Mean population size of the best 1,000 simulations over past 450,000 years.
